# An auto-generated ADP-ribosyl-cysteine degron drives DTX2-dependent turnover of PARP7

**DOI:** 10.64898/2026.10.26.761924

**Authors:** Chunsong Yang, Krzysztof Wierbiłowicz, Hyukjin Jang, Joshua S Wu, Sara C Buch-Larsen, Grant R Diefenderfer, Thomas L Genetta, Jesper V Olsen, James M Larner, Michael L Nielsen, Aakrosh Ratan, Bryce M Paschal

## Abstract

The mono-ADP-ribosyltransferase PARP7 exerts negative feedback on androgen receptor (AR) signaling by generating an ADP-ribosyl-cysteine degron on AR that marks it for degradation. Here we show that the protein half-life of PARP7 is set by cysteine auto-ADP-ribosylation, providing a mechanism for temporal control of its feedback on AR-dependent transcription in prostate cancer cells. ADP-ribosyl-cysteine in the PARP7 catalytic domain constitutes a degron read by the E3 ligase DTX2, driving PARP7 polyubiquitination and proteasomal degradation. Thus, the same E3 is used to degrade both PARP7 and its substrate AR. Using label-free mass spectrometry, we identify the AP-1 factor c-JUN as a PARP7 substrate and show that ADP-ribosylated c-JUN accumulates upon DTX2 depletion. ADP-ribosyl degron formation is dependent on NAD^+^ availability since depleting NAD^+^ pharmacologically or under hypoxia prevents degron formation and stabilizes PARP7 in a catalytically constrained state that compromises its feedback activity. Correlations between PARP7 expression and hypoxia signatures in prostate tumors, together with sensitivity to the PARP7-selective inhibitor RBN-2397 across cancer lines, nominate hypoxic, NAD^+^ depleted contexts as heightened vulnerability to PARP7-targeting agents. Marking transcription factors for degradation by covalent ADP-ribosylation is emerging as a distinct mode of transcriptional control.

## INTRODUCTION

Androgen signaling through the androgen receptor (AR) drives prostate cancer growth via ligand-dependent transcriptional programs that require termination mechanisms to limit gene activation. We recently showed that termination of androgen signaling can be mediated by an AR degradation mechanism based on the post-translational modification, mono-ADP-ribosylation. This multi-step pathway involves the mono-ADP-ribosyltransferase PARP7, which exerts negative feedback on androgen signaling by modifying cysteine residues in the AR DNA-binding domain (Wierbiłowicz *et al*, 2025). The ADP-ribosyl mark generated by PARP7 is read by an E3 ubiquitin ligase, DTX2 (Yang *et al*, 2021). Recognition of AR-conjugated ADP-ribose by the C-terminal RING-DTC (RD) domain of DTX2 drives AR ubiquitylation and proteasomal degradation. The pathway broadly parallels the mechanism of phosphor-degrons and other established post-translational modification-based turnover pathways but relies on a chemically distinct conjugate generated by a PARP enzyme (Ahmed *et al*, 2020; Wierbiłowicz *et al*., 2025). ADP-ribosylated AR can also be read as part of a non-degradative ubiquitin ligase pathway by the heterodimeric DTX3L/PARP9 complex. In this case, PARP9 supplies tandem macrodomains that directly bind ADP-ribosyl-cysteines and DTX3L contributes RING-dependent ubiquitin ligase activity (Yang *et al*., 2021), though the recruitment of this complex is not linked to AR degradation. Rather, DTX3L E3 activity modulates a subset of AR target genes likely through mono-ubiquitylation of chromatin substrates (Yang *et al*., 2021). These features illustrate how a chemical mark written by a single mono-ADP-ribosyltransferase can produce different outcomes depending on which E3 is recruited.

In addition to promoting degradation of substrates, work from multiple groups indicates that PARP7 promotes self-degradation, resulting in extraordinarily rapid turnover. In prostate cancer cells, PARP7 protein has a half-life of approximately 5 minutes), dramatically shorter than that of other PARP family members (Kamata *et al*, 2021); this can be extended by blocking catalytic function either by mutation or the selective inhibitor RBN-2397. The dependence of PARP7 degradation on its own catalytic function is notable. If PARP7 activity is linked to its turnover, this implies the presence of an auto-regulatory mechanism in which PARP7 modifies itself in addition to substrates. Such a mechanism would couple PARP7 protein abundance to the duration of PARP7 catalytic output towards substrates. In the case of transcription factors, this feature could limit the duration of negative feedback imparted by PARP7.

Support for such an auto-regulatory mechanism comes from another gene expression pathway controlled by PARP7. PARP7 was originally identified as TiPARP, the dioxin-inducible PARP, and it represses aryl hydrocarbon receptor (AHR) transactivation as part of a negative feedback loop induced by AHR’s own ligand-activated transcriptional program (MacPherson *et al*, 2013). In this context, the Ahel laboratory recently used a ubiquitin pathway blockade strategy to measure endogenous ADP-ribosylation at physiological expression levels, and showed that combined AHR activation and proteasome inhibition results in robust ADP-ribosylation of both PARP7 and AHR. They identified DTX2 as the E3 ligase that degrades both, establishing that PARP7-DTX2 crosstalk enables rapid shutdown of AHR-driven transcription (Gorelik *et al*, 2026). The biochemical basis of this mechanism, including the acceptor sites on PARP7, the sequence requirements for degron formation, and how DTX2 recognition is converted into productive polyubiquitination, remained open questions.

The DTX family of E3 ligases provides the structural basis for coupling ADP-ribose substrate recognition to ubiquitin transfer from an E2. The DTX2 RD domain harbors an ADP-ribose-binding pocket that recruits ADP-ribosylated proteins for ubiquitination, a property conserved across the DELTEX family (Ahmed *et al*., 2020). A related ADP-ribosyl-ubiquitin modification is generated during the DNA damage response (Chatrin *et al*, 2026), possibly to target a subset of proteins for degradation. How DTX2 selectively engages ADP-ribosylated degrons rather than the broader pool of ADP-ribosylated proteins, and how this recognition is converted into processive polyubiquitination, remains unclear.

NAD^+^ is the co-substrate for ADP-ribose transfer, and its availability fluctuates across physiological and pathological states including DNA damage, hypoxia, and metabolic stress. Cancer cells maintain elevated NAD^+^ through NAMPT-driven salvage synthesis to support proliferative and DNA-repair demands, and NAMPT overexpression in prostate cancer correlates with disease progression (Garten *et al*, 2009; Wang *et al*, 2011). The extent to which the PARP7-DTX2 degradation axis defined for AR and AHR signaling is sensitive to NAD^+^ availability, and whether tumor microenvironmental conditions that limit NAD^+^ alter PARP7 abundance and activity in ways relevant to therapeutic targeting, are important questions.

Here we establish that PARP7 undergoes auto-ADP-ribosylation on cysteine residues, including sites within its catalytic domain, generating a degron read by the same E3 ligase, DTX2, that recognizes ADP-ribosylated AR. We suggest that Cys543 and Cys552 are the principal sites for degron-formation by auto-modification, and show that DTX2 recognizes ADP-ribosyl-cysteine through its RD domain to drive PARP7 ubiquitylation and degradation. We provide evidence that the PARP7-DTX2 mechanism likely extends to the AP-1 factor c-JUN, which we identify as a new PARP7 substrate. We further show that the PARP7-DTX2 pathway is governed by cellular NAD^+^ availability, linking PARP7 protein turnover to metabolic state, and that hypoxic conditions characteristic of the tumor microenvironment stabilize PARP7 by limiting NAD^+^-dependent auto-modification and turnover. Together with data from other groups, our findings define a self-limiting biochemical circuit in which PARP7 catalytic output specifies its own degradation, helping shape the duration and magnitude of certain AR and AHR target gene responses in prostate cancer.

## RESULTS

PARP7 mono-ADP-ribosylates the androgen receptor (AR), generating a degron that targets AR for proteasomal degradation (Wierbiłowicz *et al*., 2025). Because PARP7 expression is induced by AR (Yang *et al*., 2021), this establishes a negative feedback loop to limit androgen-regulated gene expression. We explored whether PARP7 uses a similar degron-based mechanism that couples catalytic activity to its own degradation. To examine if PARP7 auto-modification generates a degron, we first asked whether proteasome inhibition with bortezomib affects the cellular level of ADP-ribosylated PARP7, which, based on the degron model, should accumulate as an intermediate in its degradation pathway (Fig. 1A). Following PARP7 gene induction by either AR or AHR (using the synthetic androgen R1881 or AHR agonist 10-chloro-7H-benzimidazo[2,1-a]benzo[de]isoquinolin-7-one; abbreviated BBQ), co-treatment with bortezomib produced a striking accumulation of ADP-ribosylated PARP7 (Fig. 1B, lanes 5 and 8). Co-treatment with the selective PARP7 inhibitor RBN-2397 stabilized PARP7, but, interestingly, blocking catalytic function and also prevented formation and accumulation of its ADP-ribosylated form (Fig. 1B, lanes 6 and 9). Given the selectivity of RBN-2397 across the PARP family and its inhibition of purified PARP7 auto-modification (Supplementary Fig. 1A), the most parsimonious explanation for these data is that auto-modification generates the form destined for degradation, and blocking catalytic function prevents its formation. Consistent with this view, ADP-ribose was detected on PARP7 immunoprecipitated from bortezomib-treated cells with the AF1521 macrodomain probe, as well as with an antibody specific for mono-ADP-ribosylation (Fig. 1C). Based on the chemical sensitivity of the linkage to HgCl_2_ treatment, which efficiently releases ADP-ribose from PARP7 as revealed by probing with AF1521 (Fig. 1D), the bortezomib-stabilized form of PARP7 is modified primarily (or exclusively) on cysteine residues.

**Fig. 1.**
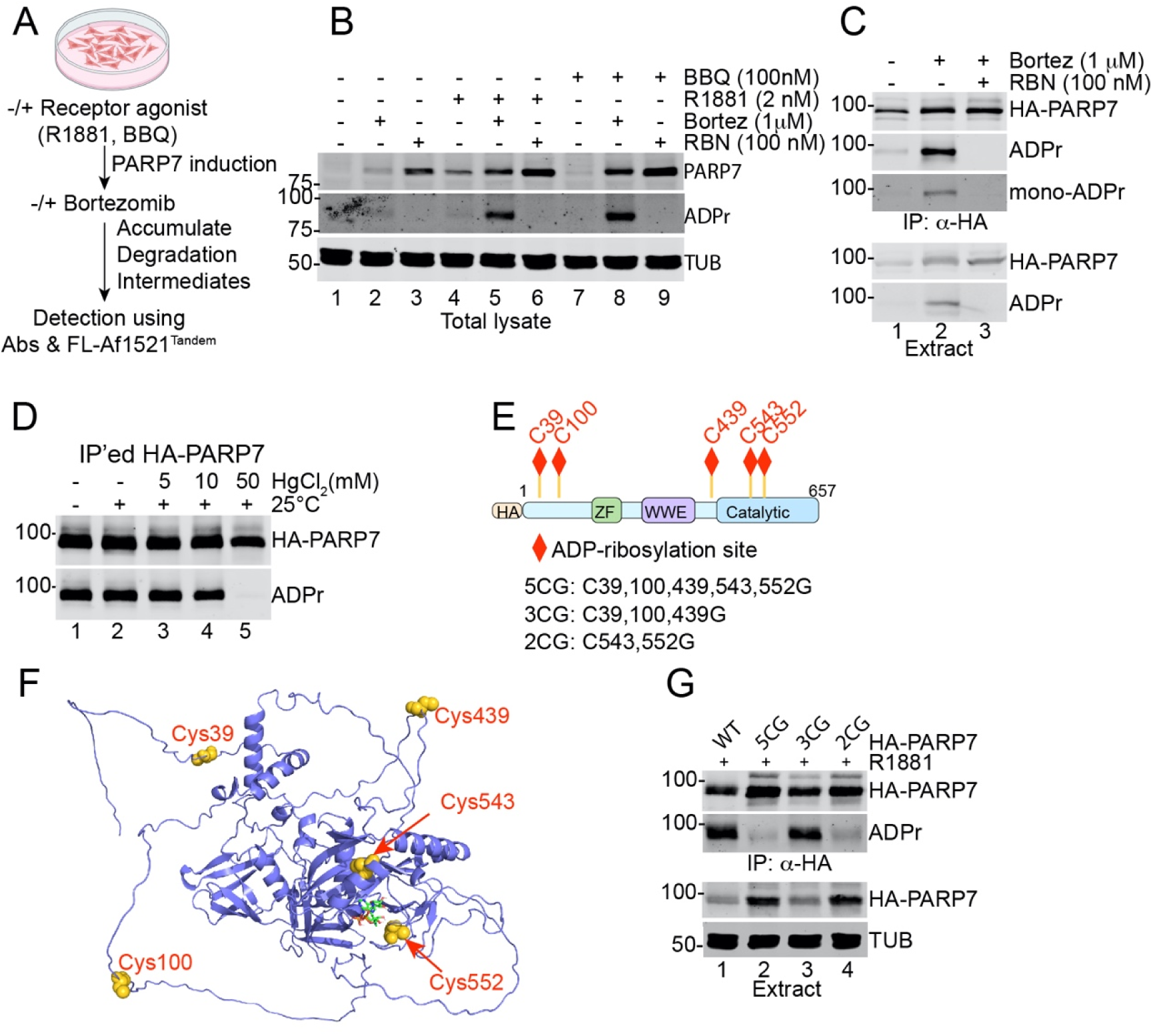
Cysteine auto-ADP-ribosylation regulates PARP7 turnover in prostate cancer cells. **A.** Workflow for PARP7 gene induction and detection of ADP-ribosyl-PARP7 in response to proteasome inhibition with bortezomib. Cell lines/drug treatments are mentioned in the methods/cell lines section. **B.** ADP-ribosylated PARP7 accumulation in cells treated with combinations of androgen (R1881), AHR agonist 10-CL-BBQ (BBQ) and bortezomib (Bort), or inhibited by PARP7 inhibitor RBN2397. Cell drug treatments were for 4 hrs. **C.** ADP-ribose detection on PARP7 isolated by IP. ADP-ribose was detected using AF1521 probe (ADPr) and via antibody (mono-ADPr). The ADP-ribose signal was eliminated by co-treatment with RBN2397. Cell drug treatments were for 4 hrs. **D.** Treating PARP7-ADP-ribose with mercury chloride releases ADP-ribose, indicating a cysteine-based linkage. Cells were treated with Bortezomib for 4 hrs, and the IP’d PARP7 was treated with HgCl_2_ at room temperature for 60 min. **E.** Cartoon of PARP7 showing its domain organization, ADP-ribosyl-Cys site locations, and the Cys-to-Gly substitutions in the PARP7 multisite mutants. **F.** Alpha-fold model of human PARP7 showing the locations of Cys acceptor sites (yellow space fill) and NAD^+^ (wireframe) in the catalytic domain as determined using Auto-Dock Vina. **G.** Detection of ADP-ribose using IP’d WT and multi-site mutants of PARP7.

### ADP-ribosylation of PARP7 on cysteines

Our group (Yang *et al*., 2021) and others (Gomez *et al*, 2018) (Rodriguez *et al*, 2021) have used mass spectrometry to identify a total of five cysteine ADP-ribosylation sites on PARP7, distributed across several of its domains (Fig. 1E). The Alpha-Fold model of human PARP7 places several of the ribosylation sites on large unstructured loops (Fig. 1F). To determine which cysteines are acceptor sites for ADP-ribose accumulation in response to bortezomib, we engineered a series of glycine substitutions (which are not acceptors for ADP-ribose) and generated stable cell lines expressing the mutant proteins. Substitution of all five cysteine sites with glycine (5CG) virtually eliminated PARP7 ADP-ribosylation, indicating that these five residues account for the bulk of auto-modification detected with AF1521 (Fig. 1G). Mutation of three cysteines located in unstructured loops (Cys39, Cys100, Cys439) had little effect on the level of PARP7 ADP-ribosylation detected on blots (Fig. 1G). By contrast, ADP-ribosylation was markedly reduced by mutation of two cysteines located in a short alpha-helix (Cys543) and an adjacent loop (Cys552) within the catalytic domain. These residues may be primary sites of PARP7 auto-modification, but we cannot rule out the possibility that amino acid substitutions at these positions reduce catalytic activity. To address the contribution of these cysteine sites for PARP7 ADP-ribosylation of substrate, we examined AR ADP-ribosylation in cells stably expressing each PARP7 mutant (Supplementary Fig. 1B). AR ADP-ribosylation by the 3CG mutant was comparable to wild-type PARP7. Substrate modification by the 2CG mutant was approximately half that of wild-type PARP7, while the 5CG mutant PARP7 was essentially inactive after correcting for the low level of AR ADP-ribosylation mediated by endogenous PARP7 in R1881-treated cells (Supplementary Fig. 1C). Thus, the 2CG mutant retains a nominal level of catalytic activity.

### Cysteine modification is linked to protein half-life

We next examined the protein half-life of the PARP7 mutants by cycloheximide chase analysis. The 3CG mutant half-life was slightly longer than wild-type protein (8.3 vs. 5.8 min; Fig. 2A, B, E). By contrast, the 5CG and 2CG mutants were substantially more stable than wild-type PARP7 (half-lives of 41 and 39 min, respectively; Fig. 2C, D, E), which is comparable to the half-life of wild-type PARP7 in RBN-2397-treated prostate cancer cells (Yang *et al*, 2023). We conclude that the three auto-modified cysteine sites in the unstructured loops of the N-terminal domain (Cys39, Cys100) and in the loop between the WWE and catalytic domains (Cys439) do not play significant roles in ADP-ribosyl-linked PARP7 degradation. By contrast, mutation of Cys543 and Cys552 in the catalytic domain promotes PARP7 stabilization, though the relative contributions of simple loss of degron formation at these sites versus impaired catalytic activity cannot be clearly distinguished by mutagenesis.

**Fig. 2.**
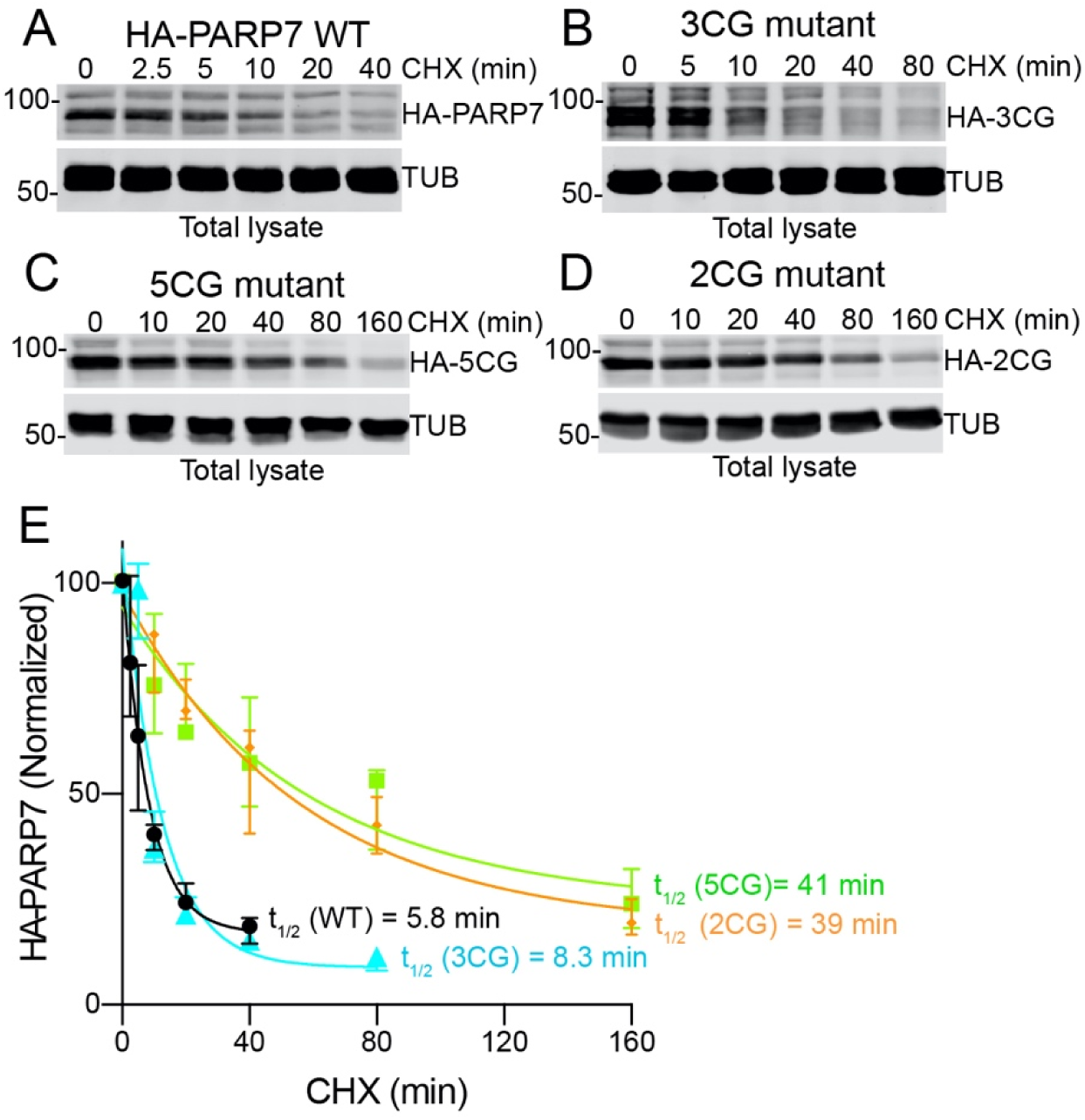
PARP7 protein half-life is dependent on ADP-ribose acceptor cysteines in the catalytic domain. **A-D.** Representative figures of cycloheximide chase analysis and immunoblotting of HA-PARP7 stably expressed in PC3-AR cell lines. **E.** HA-PARP7 blotting data (from panels A-D and Supplementary Fig. 1D) plotted to determine half-life values. The plots were generated using biological triplicate. Error bars represent standard deviation.

### E3 recognition of ADP-ribosyl cysteines

We reasoned that PARP7 auto-modification could generate a degron recognized by an E3 ubiquitin ligase harboring an ADP-ribose reader domain, as described for AR (Wierbiłowicz *et al*., 2025) and AHR (Gorelik *et al*., 2026). To test for such a mechanism, we used depletion and gene knockout approaches to evaluate E3 ligases with ADP-ribose reader domains (Fig. 3A), as well as the E3 ligases SPOP and UBR5, which lack reader domains but are known to regulate hormone signaling pathways. We predicted that depletion of an E3 that facilitates degradation of ADP-ribosylated PARP7 would result in accumulation of the modified form, detectable on blots with AF1521 probe. Of the nine E3 ligases we tested in prostate cancer cells, only DTX2 depletion resulted in accumulation of ADP-ribosyl-PARP7 (Fig. 3A, lane 3; Supplementary Fig. 2). Multiple independent siRNAs confirmed that a small reduction in DTX2 expression is sufficient to increase ADP-ribosyl-PARP7 levels (Fig. 3B; Supplementary Fig. 2). From these data we conclude that DTX2 is the major E3 ligase for PARP7 in PC3 prostate cancer cells. As an orthogonal approach, we used cell extracts from siRNA-treated PC3 prostate cancer cells for pulldown assays on AF1521 beads (Fig. 3C). Recovery of PARP7 on AF1521 beads was enriched after DTX2 knockdown, reflecting the capture of ADP-ribosylated PARP7 prior to its ubiquitylation. DTX2 was independently shown by the Ahel group to recognize and degrade ADP-ribosyl-PARP7 in HCC4 and A549 cells (Gorelik *et al*., 2026), indicating the DTX2-PARP7 degradation mechanism operates across multiple cancer cell types.

**Fig. 3.**
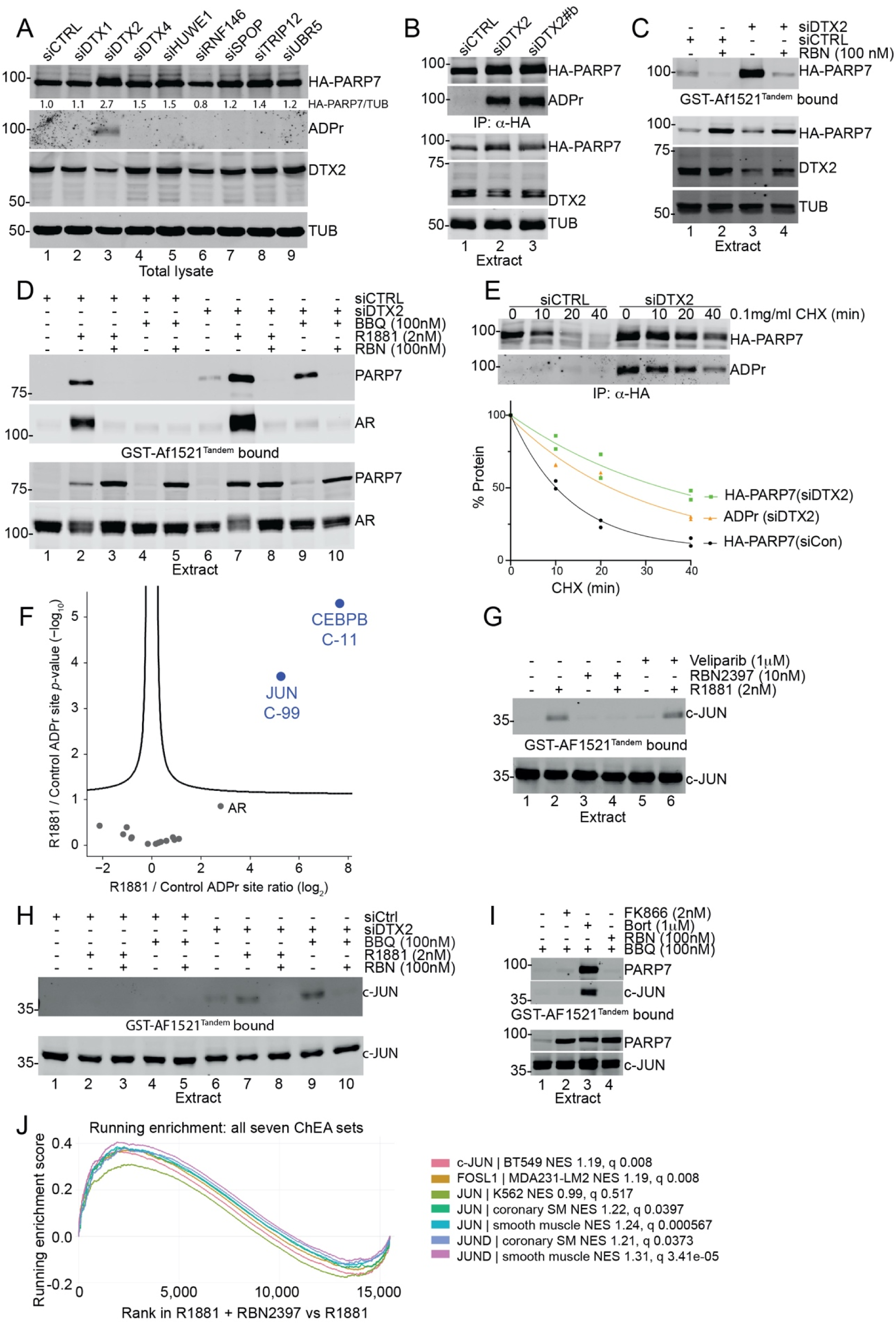
E3 ubiquitin ligase DTX2 controls the turnover of mono-ADP-ribosylated PARP7. **A.** Focused siRNA screen for E3’s that regulate the cellular level of ADP-ribosylated PARP7. **B.** Detection of ADP-ribose modification on PARP7 isolated by IP from cells depleted of DTX2. **C.** Capture of ADP-ribosylated PARP7 on AF1521 beads is sensitive to DTX2 levels and reduced by the PARP7 inhibitor RBN2397. **D.** Detection of endogenous ADP-ribosylated PARP7 by AF1521 pull-down is observed with gene induction with R1881 and is enhanced by DTX2 depletion. The level of ADP-ribosylated PARP7 induced with BBQ is below assay detection unless cells are also depleted of DTX2. **E.** ADP-ribose conjugated to PARP7 and PARP7 protein have similar turnover characteristics in DTX2 depleted cells. PC3-AR-HA-PARP7 cells treated with siCTRL and siDTX2, and HA-PARP7 and ADP-ribose were detected after IP. The experiment is done in biological duplicate. **F.** Volcano plot analysis visualizing the dynamics of ADPr sites upon treatment with R1881. Data was filtered for 4 out of 4 in at least one condition, and values imputed from normal distribution. Significance was determined via two-tailed Student’s t-testing, with an FDR of 0.05, an s0 of 0.1, and 2500 rounds of randomization. Proteins significantly upregulated by R1881 treatment are depicted in blue, and proteins not significantly regulated are shown in grey. **G.** c-JUN ADP-ribosylation in response to androgen signaling is prevented by the PARP7 inhibitor RBN2397 but unaffected by the PARP1/2 inhibitor veliparib. PC3-AR cells were treated for 4 hrs with indicated drugs and extracts subjected to pulldown on AF1521 beads. **H.** DTX2 depletion increases the level of ADP-ribosylated c-JUN detected by AF1521 pull-down in response to R1881 and BBQ treatment of PC3-AR cells. **I.** ADP-ribosylated c-JUN becomes stabilized in response to proteasome inhibition with bortezomib. c-JUN was detected by AF1521 pull-down in extracts from cells treated with the drug combinations indicated. **J.** Gene set enrichment analysis of seven ChIP-derived AP-1 transcription factor target gene sets (ChEA 2022) in VCaP cells treated with R1881 and RBN-2397 compared with R1881 alone. Running enrichment scores are plotted against genes ranked by differential expression. Normalized enrichment scores (NES) and Benjamini-Hochberg-adjusted P-values (q) are indicated for each gene set.

Several of the preceding experiments used epitope-tagged HA-PARP7 for ease of detection. To determine whether endogenous PARP7 accumulates as an ADP-ribosylated intermediate following DTX2 depletion, we induced PARP7 via AR and AHR ligands, prepared cell extracts, and performed pulldowns with AF1521 beads. ADP-ribosyl-PARP7 was recovered on AF1521 beads after gene induction; recovery was increased by DTX2 depletion and abolished by co-treatment with RBN-2397 (Fig. 3D; lanes 2, 7, 8 and lanes 4, 9, 10). DTX2 depletion also increased PARP7 protein half-life (Fig. 3E). In cycloheximide chase experiments, HA-PARP7 protein and ADP-ribosyl-PARP7 detection showed similar kinetics, consistent with a model by which the ADP-ribosyl mark is linked to PARP7 turnover (Fig. 3E).

### Cysteine ADP-ribosylation of transcription factors

Multiple transcription factors have been reported as PARP7 substrates, though AR is the only one characterized thus far in prostate cancer cells (Yang *et al*., 2021). To ask whether additional transcription factors are modified by PARP7, and whether the ADP-ribosylated forms are targets of DTX2, we set out to identify new PARP7 substrates in prostate cancer cells using AF1521 peptide enrichment coupled with label-free mass spectrometry (Supplementary Fig. 3A). Because PARP7 in prostate cancer cells is induced by androgen, the mass spectrometric analysis focused on identifying ADP-ribosylated targets upon androgen treatment. In total we identified 55 localized ADP-ribose sites, with the majority of sites occurring on serine residues followed by cysteine residues (Supplementary Table). Volcano plot analysis identified two transcription factors, c-JUN and C/EBPβ, as the only significantly enriched ADPr sites upon androgen treatment (Fig. 3F). Annotated MS/MS spectra confirmed are ADP-ribosylation on Cys99 of c-JUN and Cys11 of C/EBPβ, respectively (Supplementary 3B-D). We focused on c-JUN since C/EBPβ was recently characterized as a PARP7 substrate (Stokes *et al*, 2026). Recovery of c-JUN on AF1521 beads required androgen treatment for PARP7 induction, was prevented by co-treating cells with RBN-2397 and was unaffected by PARP1 inhibition (Fig. 3G). These results suggest that c-JUN ADP-ribosylation is mediated by PARP7. DTX2 depletion increased c-JUN recovery on AF1521 beads, and this was enhanced when PARP7 was induced by R1881 or the AHR ligand BBQ (Fig. 3H, lanes 1, 6, 7 and 9), indicating dependence on PARP7 catalytic activity. Bortezomib treatment stabilized the ADP-ribosylated form of c-JUN, similar to the effect observed for PARP7 (Fig. 3I) and AR (Wierbiłowicz *et al*., 2025). To determine whether PARP7 catalytic activity affects AP-1-dependent transcription, we analyzed RNA-seq data from androgen-stimulated VCaP cells treated with or without RBN-2397. Gene set enrichment analysis (GSEA) using seven published ChIP-derived AP-1 target gene sets identified positive enrichment in all seven sets, with six reaching statistical significance (Fig. 3J; adjusted P < 0.05). These results are consistent with PARP7-mediated repression of AP-1-dependent transcription and suggest that PARP7-mediated ADP-ribosylation and DTX2-dependent degradation extend beyond ligand-regulated transcription factors to include the signal-responsive factor c-JUN.

Notably, c-JUN represents the second AP-1 family transcription factor shown to be ADP-ribosylated on cysteine by PARP7, given the identification of Cys97 in FRA1 as a PARP7 modification site (Manetsch *et al*, 2023). We explored whether there is an enzyme-substrate gene expression relationship detectable as tumor mRNA co-expression. PARP7 mRNA correlated with both JUN (ρ=0.47, n=498, p<2.2×10 ¹) and FOSL1 (ρ=0.41, n=498, p<2.2×10 ¹) (Supplementary Fig. 3E-F), consistent with shared transcriptional regulation of PARP7 and its AP-1 substrates. Biochemically, PARP7 ADP-ribosylation has opposing consequences for these substrates. FOSL1 activity is positively regulated, while the E3 ligase DTX2 promotes degradation of ADP-ribosylated c-JUN through a mechanism similar to that targeting auto-ADP-ribosylated PARP7 itself.

### ADP-ribosyl-PARP7 interaction with DTX2

To examine the interaction between PARP7 and DTX2 in cells, we performed co-immunoprecipitation experiments under conditions that either stabilize ADP-ribosyl-PARP7 (bortezomib) or prevent its formation (bortezomib + RBN-2397). Bortezomib treatment produced substantial DTX2 enrichment in the PARP7 co-IP, which was largely eliminated by co-treatment with RBN-2397 (Fig. 4A, lanes 3, 6 and 7; Supplementary Fig. 4). We noted that HA-PARP7 co-transfection reduced Flag-DTX2 protein levels in a manner blocked by RBN-2397; whether this reflects an effect on DTX2 stability or an outcome related to transient transfection is unclear. Regardless, the co-IP data indicate that DTX2 interaction is specific for the ADP-ribosylated form of PARP7. Based on structural and biochemical studies from the Ahel and Huang groups (Ahmed *et al*., 2020; Zhu *et al*, 2022), this interaction is predicted to reflect DTX2 RD domain engagement with ADP-ribose on PARP7. We tested this prediction using cell extracts containing wild-type or cysteine mutant PARP7 in pulldown assays with recombinant GST-DTX2 RD domain immobilized on beads. Wild-type PARP7 recovery on beads was low under basal conditions but was enhanced by bortezomib treatment and abolished by co-treatment with RBN-2397 (Fig. 4B, lanes 1-3). Comparable results were obtained with the 3CG mutant, indicating that RD binding does not require ADP-ribosylation of Cys39, Cys100, and Cys439 (Fig. 4B, lanes 4-6). Only background levels of the 2CG mutant were recovered, indicating that the RD domain either does not engage these sites or that the level of auto-modification is too low for capture on beads.

**Fig. 4.**
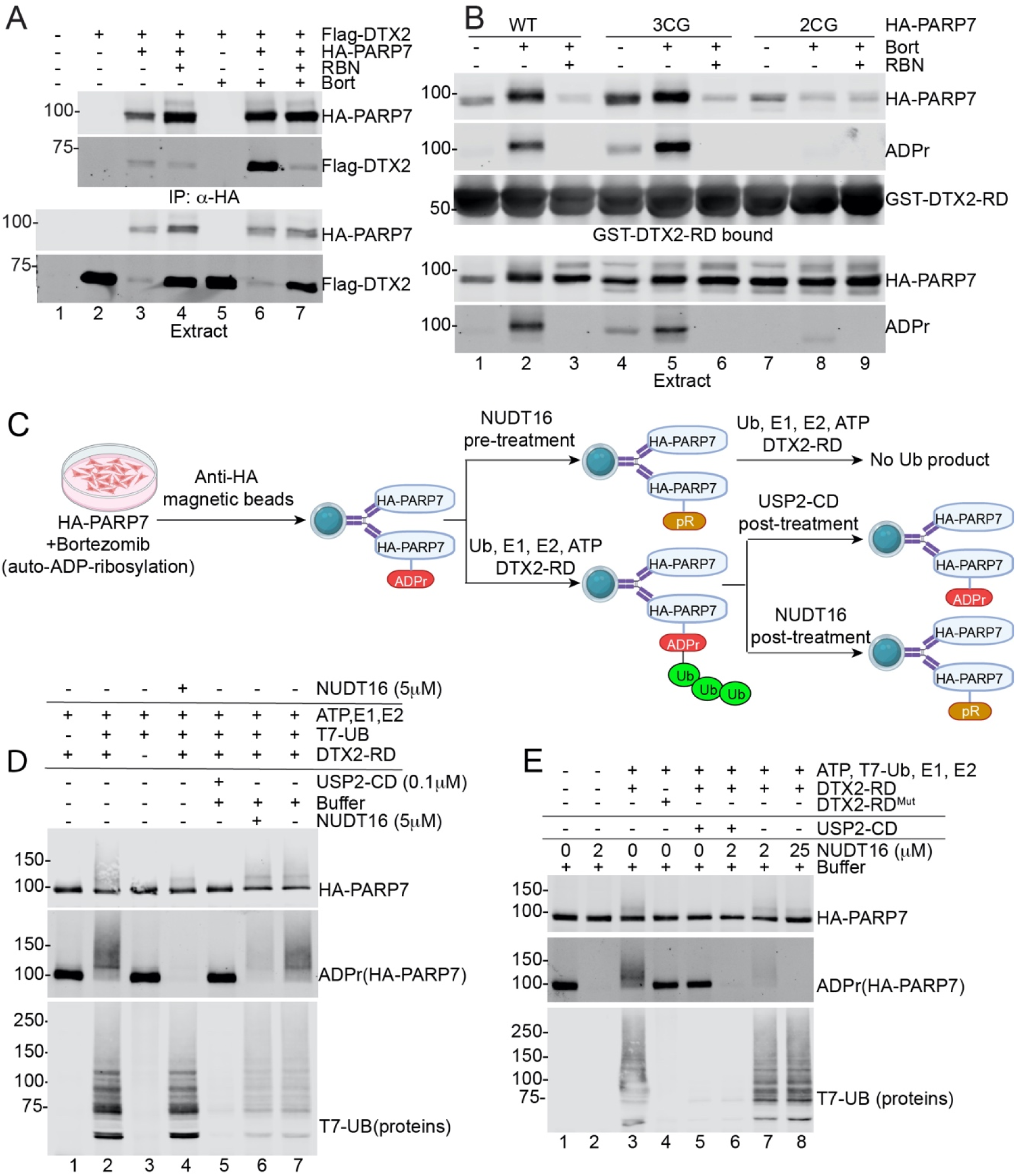
The E3 DTX2 uses an ADP-ribose reader domain to recognize the ADP-ribose degron in PARP7. **A.** Co-IP of Flag-DTX2 and HA-PARP7 depends on PARP7 ADP-ribosylation. The interaction is enhanced by bortezomib and prevented by RBN2397. **B.** Pull-down assays using recombinant GST-DTX2-RD and extracts from WT and mutant HA-PARP7 expressing cells treated with bortezomib and RBN2397. Bortezomib increased the binding of ADP-ribosylated PARP7 to DTX2 and the effect was independent of Cys39, Cys100, and Cys439, but dependent on Cys543 and Cys552 within the catalytic domain. **C.** Schemes for the ubiquitylation assays. ADP-ribosylation of HA-PARP7 was accumulated by treating the cells with 1 μM Bortezomib. ADPr-HA-PARP7 was IP’d and used as ubiquitylation substrates of DTX2 to produce Ub-ADPr-HA-PARP7. NUDT16-pretreatment of ADPr-HA-PARP7 converts it to phosphor-ribosyl (pR)-HA-PARP7 which is not a ubiquitylation substrate. USP2 post-treatment of Ub-ADPr-HA-PARP7 removes ubiquitin from ADPr and exposes ADPr for detection. NUDT16 post-treatment of Ub-ADPr-HA-PARP7 converts it to pR-HA-PARP7 which will not be detected by ubiquitin antibody or ADPr probe. **D.** ADPr-HA-PARP7 ubiquitylation by GST-DTX2-RD, with NUDT16 pre-treatment or USP2-CD and NUDT16 post-treatment. **E.** Amino acid substitutions of DTX2 RD domain that impair ADP-ribose recognition eliminate its activity as an E3 for ADP-ribosylated PARP7.

We performed in vitro ubiquitylation assays to test whether ADP-ribose conjugated to cysteine sites in PARP7 constitutes a degron that can be conjugated with ubiquitin (Fig. 4C). Using ADP-ribosyl-PARP7 immunoprecipitated from bortezomib-treated cells as substrate, polyubiquitylation of PARP7 was strictly DTX2-dependent, and the high-molecular-weight ubiquitylated products were not generated if ADP-ribose was cleaved by NUDT16 during a pre-treatment (Fig. 4D, lanes 1-4). Post-treatment with recombinant USP2 catalytic domain collapsed the ubiquitin smear to a single PARP7 band, confirming that the high-molecular-weight species reflect polyubiquitylated PARP7 (Fig. 4D, lanes 2 and 5). Post-treatment with NUDT16 reduced the level of Ub-ADP-ribosyl-PARP7 detected by the AF1521 probe without altering the overall ubiquitin pattern detected with T7-Ub (Fig. 4D, lanes 6 and 7). To confirm that DTX2-mediated ubiquitylation requires RD domain recognition of ADP-ribose, we compared wild-type RD to a mutant bearing three substitutions in the ADP-ribose binding pocket. The DTX2 RD domain mutant failed to support ubiquitin addition to ADP-ribosyl-PARP7 (Fig. 4E, lanes 3 and 4). We also found that a higher concentration of NUDT16 (25 μM) was more effective at cleaving ADP-ribosyl smears detected by AF1521 (Fig. 4E, lanes 7 and 8). We speculate that the relative resistance of poly-Ub-ADP-ribosyl-PARP7 to NUDT16 could reflect steric protection of the ADP-ribose linkage that helps protect the conjugate prior to proteasomal degradation.

### NAD^+^-sensitive PARP7 regulation

Because DTX2-dependent ubiquitylation is strictly dependent on the ADP-ribosyl degron generated by PARP7 auto-modification, we predicted that this mechanism would be sensitive to cellular NAD^+^ levels. The Kraus laboratory recently showed that during early adipocyte differentiation, a decline in NAD^+^ levels coincides with increased PARP7 expression at both the RNA and protein levels, and that NMN supplementation increased ADP-ribose on immunoprecipitated PARP7 while reducing PARP7 protein abundance (Stokes *et al*., 2026). These data were interpreted as evidence that PARP7 senses transient NAD^+^ fluctuations during cell differentiation. In cancer cells, NAD^+^ is maintained at high levels by overexpression of nicotinamide phosphoribosyltransferase (NAMPT), the rate-limiting enzyme of the NAD^+^ salvage pathway that helps meet the demand created by high metabolic activity, DNA damage, and constitutive PARP activation (Kennedy *et al*, 2016). In prostate cancer specifically, NAMPT overexpression correlates with disease progression and castration resistance (Wang *et al*., 2011). To test the effect of NAD^+^ levels on PARP7 stability, we employed the pharmacological NAMPT inhibitors FK866 and STF-118804. Endogenous PARP7 was induced via AHR activation with BBQ (100 nM) and cells were co-treated with FK866 (10 nM) or STF-118804 (50 nM). Immunoblotting revealed a striking accumulation of PARP7 protein under both conditions, indicating that PARP7 protein is sensitive to NAD^+^ levels (Fig. 5A). We predicted that NAD^+^ limitation would also impair PARP7-mediated negative feedback on AHR target gene expression. Consistent with this, co-treatment of PC3 cells with BBQ and FK866 produced a dramatic increase in AHR-driven induction of the PARP7 and CYP1A1 genes (Fig. 5B), paralleling results reported with RBN-2397-mediated inhibition of PARP7 catalytic activity (Stokes *et al*., 2026). Together, these data indicate that both PARP7 protein turnover and its negative feedback action on AHR target gene expression are sensitive to cellular NAD^+^ levels. The FK866 effect was rescued by supplementation with nicotinamide riboside (NR), which restores NAD^+^ levels by bypassing NAMPT via the NRK1/2-dependent pathway (Fig. 5C); this result establishes the on-target effect of FK866.

**Fig. 5.**
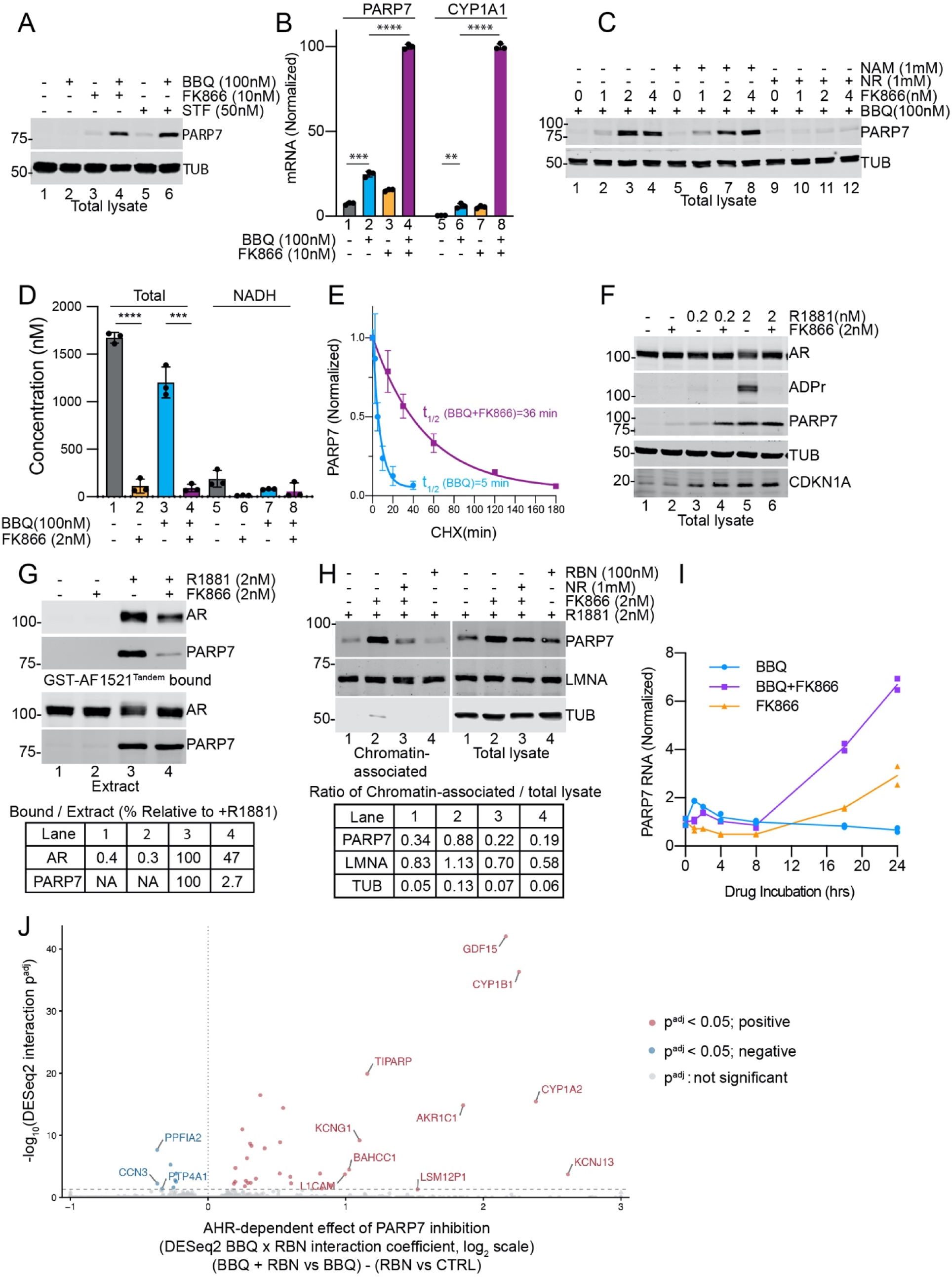
Cellular NAD^+^ levels are important for the rate of PARP7 turnover, which impacts AR and AHR-dependent gene regulation. **A.** Reducing NAD^+^ levels with selective nicotinamide phosphoribosyl-transferase (NAMPT) inhibitors increases PARP7 protein levels in cells. PC3-AR cells were treated with BBQ in the absence and presence of FK866 and STF-118804 and the levels of endogenous PARP7 were determined by immunoblotting. **B.** AHR target genes PARP7 and CYP1A mRNA levels are significantly increased by NAD^+^ depletion. The experiment is done in biological triplicate. Error bars represent standard deviations. Statistical significance P value **** <0.0001, *** <0.001, ** <0.01. **C.** Supplementation with the NAD precursor nicotinamide riboside (NR) bypasses NAMPT inhibition and restores PARP7 protein turnover. PC3-AR cells were treated with the indicated drug combinations and PARP7 protein levels determined by immunoblotting. **D.** NAD^+^ levels are reduced by treating PC3-AR cells with nanomolar levels of FK866. Total (NAD^+^ and NADH) and NADH levels in cell extracts were measured in technical triplicate and plotted. **E.** PARP7 protein half-life is increased approximately 7-fold by reducing cellular NAD^+^ levels. The assays were done in biological triplicate. Error bars represent standard deviations. **F.** Depletion of NAD^+^ by FK866 treatment stabilizes PARP7 but limits AR ADP-ribosylation. Proteins were detected by immunoblotting, and ADP-ribosylation of AR determined using an AF1521 probe. **G.** AF1521 pulldown of ADP-ribosylated AR and PARP7 from NAD^+^-depleted cells demonstrates both substrate and auto-modification are sensitive to NAD^+^ levels. **H.** PARP7 stabilized by NAD^+^ depletion partitions with the Triton X-100-resistant fraction of cells. **I.** NAD^+^ depletion causes loss of the negative feedback on AHR and hyperactivation of the AHR target gene PARP7. **J.** Volcano plot showing the interaction between AHR activation by BBQ and PARP7 inhibition by RBN-2397 in VCaP cells. The x-axis represents the interaction log fold change, and the y-axis represents −log of the adjusted P-value. Positive values indicate an increased response to RBN-2397 in the presence of BBQ relative to its effect without BBQ. Significantly altered genes (adjusted P < 0.05) are highlighted. For visualization, the 10 most significant genes with positive interaction coefficients and the 3 most significant genes with negative interaction coefficients are labeled.

We confirmed that FK866 reduces NAD^+^ levels in PC3 cells (Fig. 5D, Supplementary Fig. 5A). Cycloheximide chase experiments measuring endogenous PARP7 induced by AHR revealed a 7-fold increase in PARP7 stability upon NAD^+^ depletion, providing evidence that PARP7 half-life is sensitive to intracellular NAD^+^ levels (Fig. 5E, Supplementary Fig. 5B-C). AR undergoes androgen-dependent, multi-site ADP-ribosylation by PARP7 (Fig. 5F, lane 5; (Yang, 2021 #107)), and this modification was reduced by FK866 co-treatment (Fig. 5F, lanes 5 and 6). To increase sensitivity for ADP-ribose detection on AR and PARP7, we used AF1521 bead pulldowns followed by blot detection. Under NAD^+^-depleted conditions, ∼47% of AR and ∼2.7% of PARP7 were recoverable in the ADP-ribosylated state (Fig. 5G). We next examined how NAD^+^ limitation affects PARP7 chromatin association by biochemical fractionation. NAD^+^ depletion increased PARP7 in the Triton X-100-insoluble nuclear fraction, suggesting that failure to auto-modify and undergo degradation results in PARP7 accumulation on chromatin (Fig. 5H). Together, these results indicate that lowering cellular NAD^+^ reduces PARP7 auto-modification and slows its turnover. Despite increased PARP7 abundance, target modification is reduced since the co-substrate NAD^+^ becomes limiting.

### NAD^+^ levels impact negative feedback by PARP7

To examine temporal effects of NAD^+^ depletion on PARP7 gene induction, we co-treated PC3 cells with BBQ and FK866 and measured RNA levels at multiple timepoints (0-24 hrs). A small early peak of AHR-induced PARP7 expression was observed at 1-2 hr under both conditions (Fig. 5I, Supplementary Fig. 5D). However, beyond 8 hours, FK866 treatment produced a striking increase in PARP7 gene induction. This timepoint corresponds to the time required for FK866-mediated NAMPT inhibition to reduce NAD^+^ to levels that are limiting for AR ADP-ribosylation (Fig. 5F). We conclude that both PARP7 protein and nominal cellular NAD^+^ concentration are required to maintain negative feedback on AHR-driven PARP7 and CYP1A1 expression (Fig. 5B). Interestingly, FK866 treatment without AHR ligand addition also induced PARP7 expression, suggesting additional mechanisms may contribute to PARP7 induction when cellular NAD^+^ is limiting. To assess the broader effects of PARP7 inhibition on AHR-dependent transcription, we analyzed RNA-seq data from VCaP cells treated with the AHR agonist BBQ, RBN-2397, or their combination. Analysis of the interaction between BBQ and RBN-2397 identified 40 genes whose response to AHR activation was significantly altered by PARP7 inhibition (adjusted P < 0.05; Fig. 5J). Of these, 28 showed enhanced induction and six showed enhanced repression. These findings indicate that PARP7 catalytic activity limits the transcriptional response to AHR activation, consistent with its role in negative feedback regulation.

Elevated NAD^+^ levels in cancer cells support heightened energy metabolism and signaling functions that promote tumor growth. However, NAD^+^ levels can be acutely or chronically reduced. Acute depletion follows DNA damage, where PARP1 activation rapidly consumes NAD^+^, while chronic depletion occurs in hypoxic tumors where reduced mitochondrial activity limits NAD^+^ regeneration. To determine whether NAD^+^ reduction in hypoxic cells affects PARP7 expression and protein turnover, we incubated PC3 cells in 0.1% oxygen for up to 18 hours and examined PARP7 by immunoblotting. PARP7 was essentially undetectable under normoxic and hypoxic conditions alike, including in the presence of BBQ (Fig. 6A, lanes 1-4), consistent with its rapid turnover. Inclusion of RBN-2397 to inhibit auto-modification-linked proteasomal degradation enabled PARP7 detection under both conditions (Fig. 6A, lanes 5, 6, 11 and 12). HIF1α immunoblotting confirmed an active hypoxic response at both the 6 and 18 hr timepoints. Measurements of NAD^+^ and NADH levels confirmed that hypoxia reduced NAD^+^ levels by approximately 75% and produced a striking decrease in the NAD^+^/NADH ratio (Fig. 6B-C; Supplementary Fig. 6A). To directly assess how hypoxia-induced NAD^+^ depletion affects PARP7 turnover, we performed cycloheximide chase assays in triplicate and measured HA-PARP7 by immunoblotting (Supplementary Fig. 6B). Hypoxia extended the PARP7 half-life to 24 minutes (Fig. 6D), corroborating our FK866 results and suggesting that NAMPT inhibition and hypoxia stabilize PARP7 through an NAD^+^-dependent mechanism. Together, these findings imply that tumor microenvironmental conditions that limit NAD^+^ availability such as hypoxia simultaneously reduce PARP7 catalytic activity toward substrates while increasing PARP7 protein stability, and provide a rationale for identifying tumor contexts with heightened vulnerability to RBN-2397 based on PARP7 expression.

**Fig. 6.**
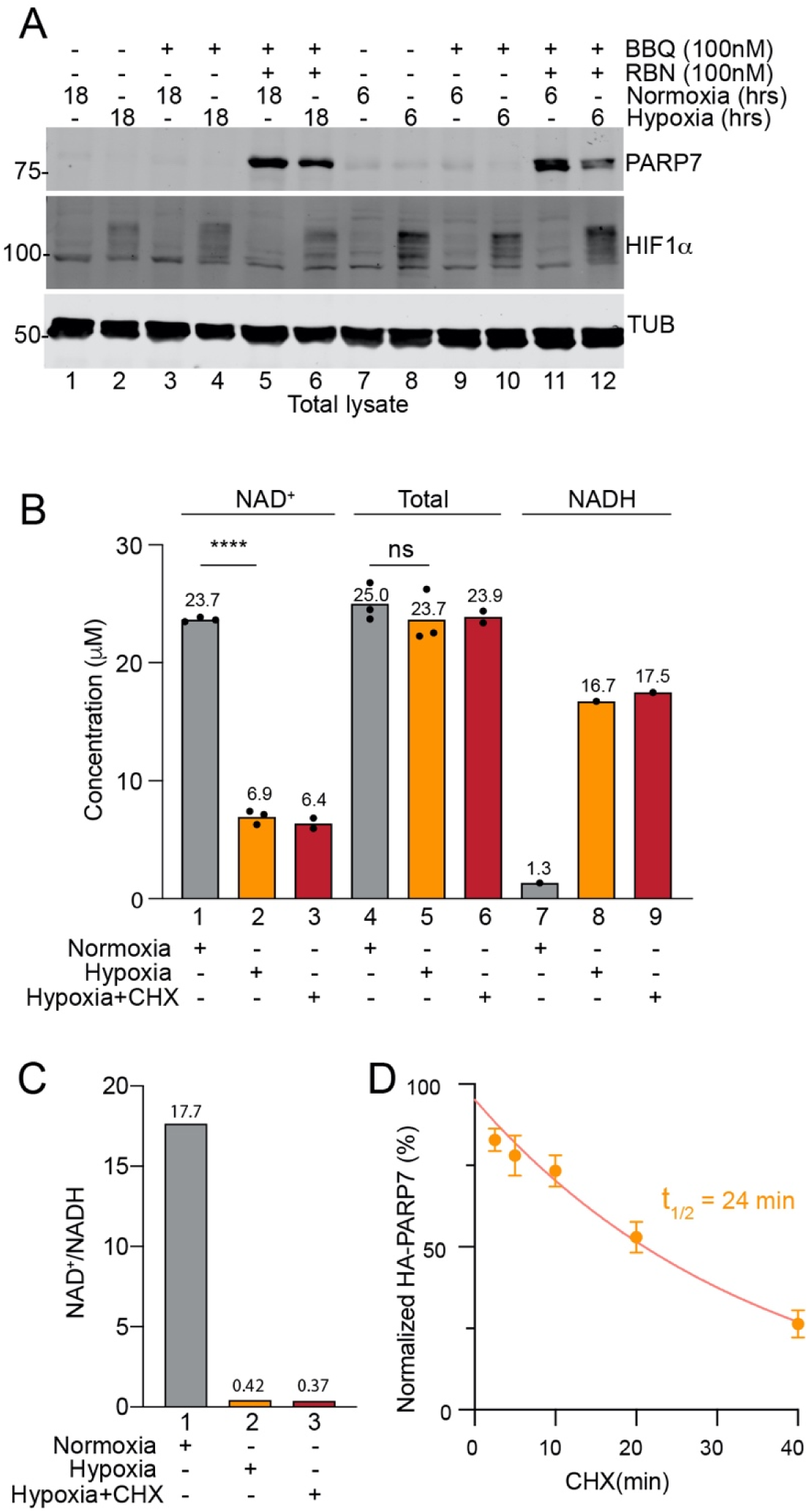
NAD^+^ depletion in hypoxia stabilizes PARP7 protein. **A.** PARP7 expression assayed by immunoblotting following overnight induction by BBQ under normoxic and hypoxic conditions. HIF1α was used as a positive control for hypoxic conditions. **B.** NAD^+^ and total (NAD^+^ + NADH) levels under normoxic and hypoxic conditions. NADH level was from total subtracted by NAD^+^ level. The dots represent technical replicate. P value **** <0.0001, ns: not significant. **C.** NAD^+^/NADH ratio measured during a cycloheximide chase assay. **D.** HA-PARP7 protein levels determined by immunoblotting and plotted as a function of time during a cycloheximide chase assay. The experiment is done in biological triplicate. Error bars represent standard deviation.

### PARP7 expression in prostate cancer

Because PARP7 induction by BBQ persists under hypoxic conditions, hypoxic tumors may represent a treatment context favorable for effective RBN2397 target engagement. We tested this prediction by analyzing the statistical relationship between RNA-seq, copy-number, and clinical data from TCGA-PRAD (localized prostate adenocarcinoma, n=497 primary tumors) and the SU2C/PCF metastatic cohort (mCRPC; n=444 metastatic samples). PARP7 mRNA correlated strongly with HIF1α mRNA (ρ=0.33, n=497, p=2.7×10□¹□) and a hypoxia gene expression signature (Hallmark Hypoxia; ρ=0.62, n=497, p<2.2×10□¹□) in TCGA-PRAD (Fig. 7A, 7B), consistent with hypoxia-driven NAD^+^ depletion increasing PARP7 expression. The presence of a hypoxia response element in the PARP7 gene implies a direct role for HIF1α in this pathway(Zhang *et al*, 2020). The relationship held after adjusting for tumor purity, Gleason grade, and AR transcriptional activity (partial ρ=0.51 to 0.56, n=333) and was uniform across the major TCGA-PRAD molecular subtypes (ERG-fusion, ETV1-fusion, SPOP-mutant, unclassified; interaction BH-adjusted p=0.83, n=303), arguing against a subtype-restricted effect (Supplementary Fig. 7).

**Fig. 7.**
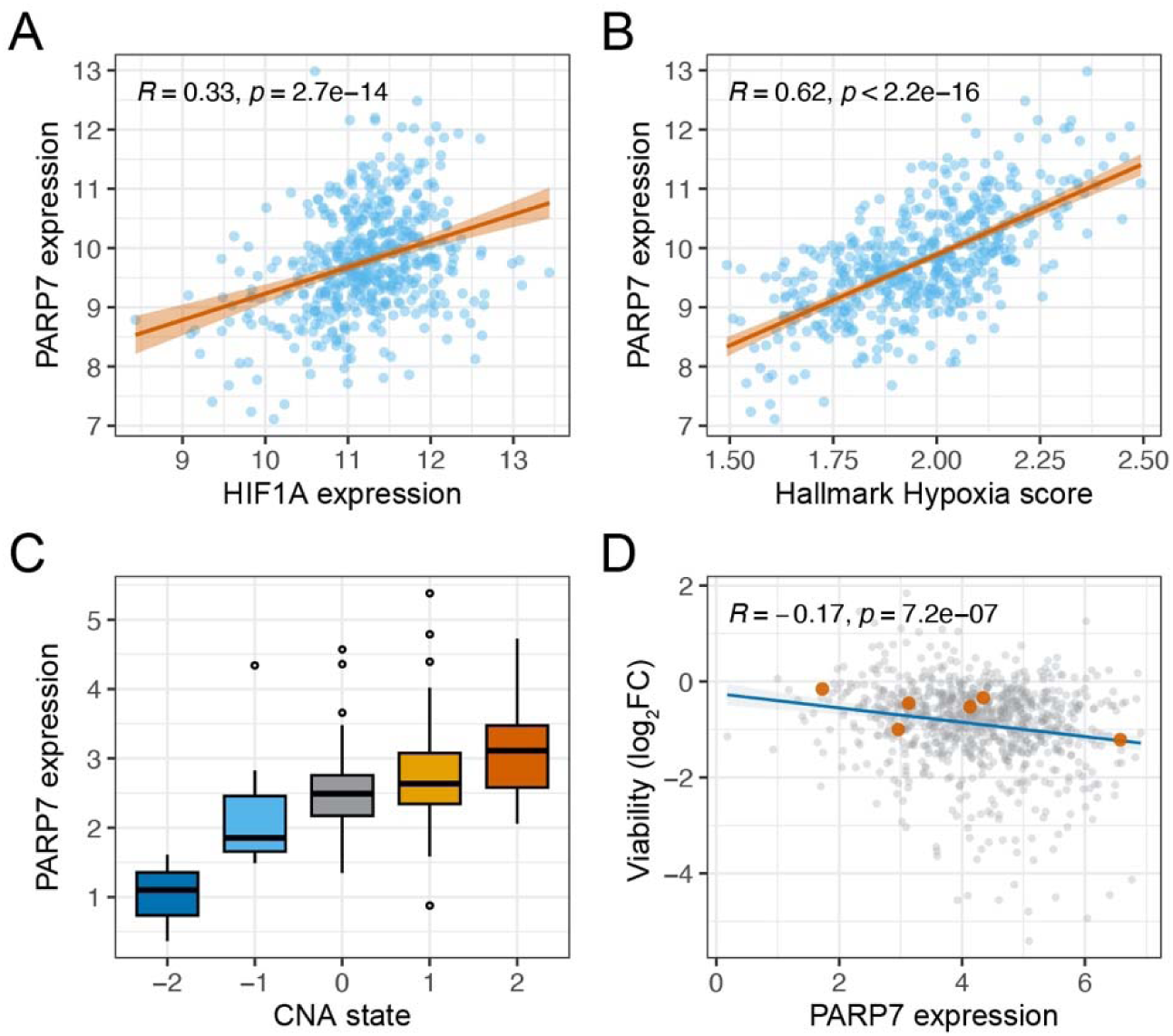
PARP7 mRNA tracks hypoxia in prostate cancer and predicts RBN2397 sensitivity. Expression values are log_2_-transformed normalized counts (TCGA-PRAD, panels A through C) or log_2_(TPM+1) (DepMap, panel D). Spearman ρ and p-value annotated on each scatter plot; regression lines with 95% confidence intervals shown. **A.** HIF1A expression versus PARP7 expression in TCGA-PRAD primary tumors (n=497). **B.** Hallmark Hypoxia ssGSEA score versus PARP7 expression in TCGA-PRAD (n=497). **C.** PARP7 expression by GISTIC2 copy-number category in the SU2C/PCF mCRPC cohort (n=262, one sample per patient). Categories used were homozygous deletion (Homdel), heterozygous loss (Hetloss), diploid, low-level gain (Gain), and high-level amplification (Amp). Ordinal Spearman correlation across the full spectrum shown. **D.** PARP7 expression (log_2_(TPM+1), DepMap 26Q1) versus RBN2397 (ATAMPARIB) viability (PRISM Repurposing Primary, log_2_ fold change; more negative indicates greater sensitivity to drug) across 859 cancer cell lines. Prostate adenocarcinoma cell lines highlighted in orange. OLS regression line shown for visual reference.

We previously reported significant differences in progression-free survival between patients stratified by PARP7 expression in TCGA-PRAD (top versus bottom quartile, n=168, 33 events), with higher PARP7 associated with improved outcome (Yang *et al*., 2021). We next asked whether the PARP7 pathway is altered in metastatic, castration-resistant disease. In the SU2C/PCF mCRPC cohort, PARP7 was focally amplified in 7.7% of tumors (34/444), and amplified tumors expressed higher PARP7 mRNA than non-amplified tumors (Wilcoxon p=0.0013, n=262), indicating that this genomic alteration is transcriptionally active. PARP7 expression increased monotonically across the copy-number spectrum (Spearman ρ=0.32, n=262, p=1.7×10□□), with low-level gains (33% of tumors) already showing elevated expression relative to diploid tumors and a further increase in amplified tumors, consistent with a graded dosage effect (Fig. 7C). The CNA-mRNA concordance and the dose-response remained significant after adjusting for biopsy site and contributing institution (n=262, adjusted p=7.0×10□□ and 1.4×10□□, respectively), arguing against a site- or batch-driven artifact. PARP7 amplification and mRNA level were not associated with overall survival, prior abiraterone/enzalutamide exposure, or neuroendocrine trans-differentiation (NEPC score), though the survival comparison was underpowered (n=80, 46 events). Together with the primary-disease mRNA correlations, these data nominate genomically amplified, transcriptionally active PARP7 as a marker of a candidate population for PARP7-directed therapy in metastatic disease, complementary to the NAD^+^/hypoxia-based rationale in primary tumors.

To test whether elevated PARP7 expression translates to RBN2397 sensitivity, we examined the Cancer Dependency Map (DepMap). Across 859 cancer cell lines with PRISM primary viability data, TIPARP expression was negatively correlated with RBN2397 viability (Spearman ρ = −0.17, n = 859, p = 7.2×10□□; Fig. 7D), indicating that cell lines with higher PARP7 expression are more sensitive to RBN2397. Together, the tumor expression data and the DepMap pharmacogenomics identify the hypoxic, NAD^+^-limited tumor microenvironment as a candidate context for RBN2397 efficacy in prostate cancer.

## DISCUSSION

Our study shows that PARP7 is degraded via an auto-generated ADP-ribosyl degron, extending the writer-reader-degradation logic previously defined for AR regulation (Wierbiłowicz et al., 2025). Proteasome inhibition in prostate cancer cells resulted in the accumulation of PARP7 ADP-ribosylated selectively on cysteine-residues; the ADP-ribosylated PARP7 degradation intermediate required PARP7 catalytic activity and was recognized by the ADP-ribose reader domain of the E3 DTX2. Mutagenesis of five cysteines known to be ADP-ribosylated points to Cys543 and Cys552 within the catalytic domain as the principal amino acids for conjugation and regulation of to PARP7 protein turnover. Three other cysteines that undergo ADP-ribosylation located in the unstructured loops are not required for the rapid turnover of PARP7 protein in cells. Thus, the degron generated by self-ADP-ribosylation is restricted to a subset of the cysteines modified by PARP7. This finding is reminiscent of AR which undergoes multi-site ADP-ribosylation in the unstructured N-terminal domain, but degron formation maps to the DNA-binding domain (Yang *et al*., 2021). Because the PARP7 Cys543,552Gly mutant (2CG) has partial catalytic activity toward AR while showing markedly extended half-life, we acknowledge that these substitutions could affect protein stability through a structural effect distinct from auto-generation of the ADP-ribosyl degron. The convergence of independent lines of evidence, reduced ADP-ribosylation signal revealed by AF1521 and antibody detection, reduced DTX2 binding in RD domain pulldown assays, and extended half-life by cycloheximide chase, together support a degron model centered on these two catalytic domain cysteines.

The identification of DTX2 as the E3 ligase that recognizes auto-ADP-ribosylated PARP7 establishes that a shared mechanism is utilized for AR and PARP7 degradation. In both cases, an ADP-ribosyl-cysteine degron is read by the DTX2 RD domain and used as a signal for proteasomal degradation. A targeted screen of E3 ligases bearing ADP-ribose reader domains here and by other groups established that DTX2 depletion causes an accumulation of ADP-ribosylated PARP7. This result corroborates and extends the key finding from the Ahel laboratory who used ubiquitin pathway blockade to show that DTX2 is the E3 ligase responsible for degrading endogenous ADP-ribosylated PARP7 and the transcription factor AHR (Gorelik et al., 2025). Their work helped to establish that PARP7 and DTX2 exert negative feedback on AHR. Our work builds on that foundation by resolving key molecular details including defining the degron-forming sites in PARP7 and linking these sites to recognition by DTX2 via its ADP-ribose-binding pocket. Further biochemical work has implied an additional E3 ligase in this pathway. It was shown that following DIX2 mono-ubiquitylation of an-ADP-ribose degron, RNF114 recognizes Ub-ADP-ribose and mediates polyubiquitylation through K11-based chain extension using the mono-ubiquitin as the seed (Kloet *et al*, 2025; Lacoursiere *et al*, 2025). Taken together, the available data suggest how three enzymes (PARP7, DTX2, RNF114) can be used in sequence to regulate protein turnover.

An additional finding of our work is that PARP7-DTX2 degron formation is not restricted to PARP7 itself but extends to c-JUN, identified through AF1521 peptide enrichment and quantitative mass spectrometry as a PARP7 substrate alongside C/EBPβ (Stokes *et al*., 2026). c-JUN ADP-ribosylation requires androgen-induced PARP7 activity, is blocked by RBN-2397 but not by PARP1/2 inhibition, increases upon DTX2 depletion, and is stabilized by proteasome inhibition with kinetics paralleling those observed for PARP7 and AR. Together with the prior identification of FRA1 Cys97 as a PARP7 acceptor site (Manetsch *et al*., 2023), c-JUN establishes a second AP-1 family member as a PARP7 substrate, and the positive mRNA co-expression observed between PARP7 and both JUN and FOSL1 in prostate tumors suggests that PARP7 and its AP-1 targets are subject to coordinated transcriptional control. Notably, the consequences of PARP7-mediated ADP-ribosylation diverge sharply between these two AP-1 substrates, since modification stabilizes FRA1 by blocking PSMC3-dependent degradation (Manetsch *et al*., 2023), whereas our data show that modification of c-JUN, like modification of PARP7 itself, creates a DTX2-dependent degron that promotes turnover. This divergence reinforces a PARP7 theme that the same biochemical mark, cysteine mono-ADP-ribosylation, can be read in different ways depending on the structural context of the modified protein and the reader engaged. Additional divergent outcome examples include PARP7 modification and stabilization of liver X receptors (Bindesboll *et al*, 2016) and destabilization of the estrogen receptor α (Rasmussen *et al*, 2021). The net transcriptional consequence of PARP7 effects on a particular transcription factor is also likely to be dependent on the cellular and tissue context, and the relative abundance and modification kinetics of the substrate.

The NAD^+^ dependence we identified converges with work from the Kraus laboratory. Studying PARP7 in adipocyte differentiation, Stokes *et al*. reported PARP7 acts as a nuclear NAD^+^ sensor, auto-modifying at high NAD^+^ levels and declining as NAD^+^ levels fall (Stokes *et al*., 2026). Pharmacologically, NAMPT inhibition (FK866, STF-118804) increased PARP7 levels, extended its half-life about 7-fold, and amplified AHR-driven PARP7/CYP1A1 induction; these effects were reversed by NAD^+^ repletion via nicotinamide riboside. NAD^+^ depletion thus increases PARP7 abundance while reducing its catalytic output. PARP7 accumulates in a catalytically constrained, unmodified state unable to modify AR or trigger its own degradation, consistent with its increased partitioning into the Triton-insoluble chromatin fraction. In hypoxia, a 75% drop in NAD^+^ levels and collapsed NAD^+^/NADH ratio extended PARP7 half-life similarly, suggesting hypoxic tumor regions sustain elevated PARP7 through impaired auto-degradation rather than transcriptional induction. These findings have direct implications for the use of RBN-2397 and related PARP7 inhibitors as cancer therapeutics, and in particular suggest that hypoxic tumors represent a context of heightened actionability for this drug class. Because PARP7 catalytic inhibition and NAD^+^ depletion both stabilize PARP7 protein by the same underlying mechanism, namely loss of auto-ADP-ribosylation-dependent degradation, the pharmacodynamic accumulation of PARP7 protein observed upon RBN-2397 treatment should not be interpreted as a marker of increased PARP7 activity; rather, it reflects the inhibitor uncoupling PARP7 abundance from PARP7 function. This distinction is important for the design of biomarker strategies and combination approaches, particularly given that PARP7 expression correlates with hypoxia signatures and HIF1A expression in TCGA prostate adenocarcinoma data, and that PARP7 expression across cancer cell lines correlates with sensitivity to RBN-2397 in pharmacogenomic screening data. The mechanism described here provides a biochemical rationale for these correlations. Hypoxic tumor regions experience chronically reduced NAD^+^ regeneration due to impaired mitochondrial function, and our data show that this NAD^+^ limitation independently elevates PARP7 protein stability by suppressing auto-degron formation, in addition to any transcriptional induction of PARP7 by HIF1α or AHR signaling (MacPherson *et al*., 2013; Zhang *et al*., 2020). Tumor cells in hypoxic niches may therefore accumulate PARP7 protein through a mechanism that is largely orthogonal to androgen or AHR ligand availability, identifying hypoxic regions of prostate tumors, and potentially other PARP7-expressing tumor types, as a population in which PARP7 protein is intrinsically stabilized and where pharmacological inhibition with RBN-2397 might achieve more durable target engagement.

## Acknowledgment

We thank Kevin Omohundro and Troy Chesser for their technical support. Funding: NCI award R01CA214872 (BMP), Saunders Memorial Funding, UVA Comprehensive Cancer Center (BMP).

## MATERIALS AND METHODS

### Cell lines

All cells were grown at 37°C in 5% CO_2_ incubator. HEK293T cells (ATCC CRL-3216, RRID CVCL_0063) were maintained in DMEM (Corning 10-017-CV) supplemented with 5% heat-inactivated fetal bovine serum (FBS, Corning 35-011-CV) and 1% penicillin-streptomycin (P/S, Cytiva SV30010). PC3 (ATCC CRL-1435), PC3M (ATCC, RRID CVCL_9555), PC3M/HA-PARP7, PC3-AR, and PC3-AR/HA-PARP7 (Yang et al, 2021) were maintained in RPMI 1640 (Corning 10-040-CV) supplemented with 5% FBS and 1% P/S. To generate PC3-AR/HA-PARP7 mutant cell lines (5CG, 3CG, and 2CG), HEK293T cells were passaged (1:3) into a 6-well plate in the morning, grown for 6 hours, and transfected with 0.9 μg of target plasmid, 0.3 μg of pMD2.G, and 0.6 μg of psPAX2 using PEI transfection reagent (MedChemExpress K-2014). After incubation at 37°C for approximately 16 hours, cells were replenished with fresh DMEM containing 35% FBS and incubated for a further 24 hours. Medium containing lentivirus was transferred to a fresh conical tube, clarified by centrifugation (approximately 700 × g), passed through a sterile 0.45 μm filter, and used to infect PC3-AR or PC3M in growth medium containing 8 μg/ml polybrene. After 2 to 3 cell doublings, cells were selected with 1 to 2 μg/ml puromycin according to the antibiotic resistance of the target plasmid. Cell lines used in each figure panel, with drug-treatment times, are listed below.

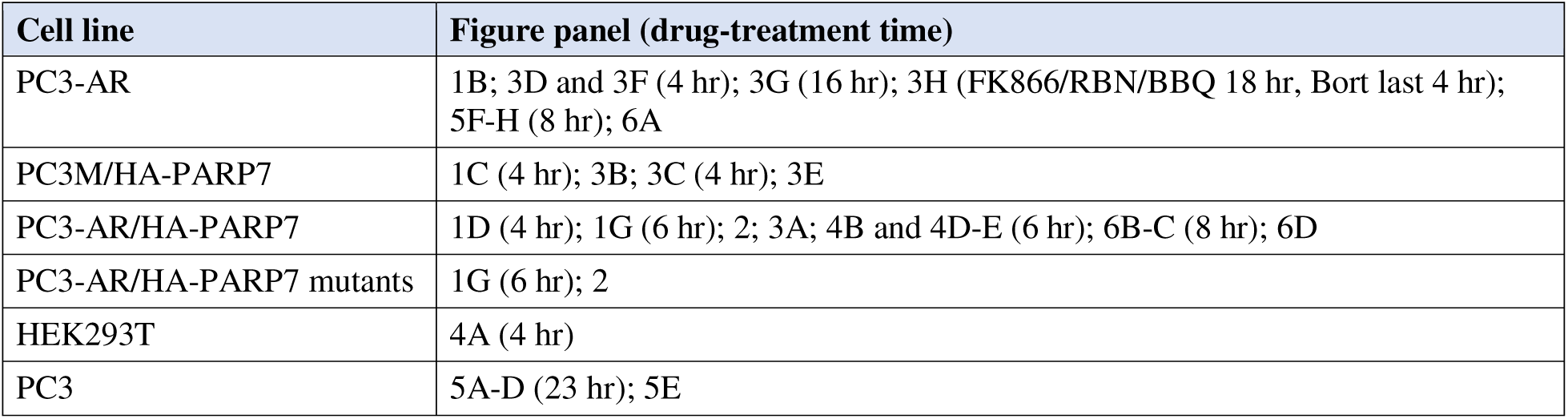

### siRNA transfection

Control siRNA (siCTRL) was from Ambion (AM4635). siRNAs targeting ubiquitin E3 ligases were from Thermo Fisher Scientific and were siDTX1 (s4356), siDTX2 (s223231), siDTX4 (s23321), siHUWE1 (s19596), siRNF146 (s37822), siSPOP (s15955), siTRIP12 (s17810), siUBR5 (s28025), siDTX2#b (s41519), and siDTX2#c (s195603). siDTX3L was custom synthesized against the target sequence CGCGTATTAGGAGTCTCAGAT. Cells were transfected with 20 nM siRNA using Lipofectamine RNAiMAX (Invitrogen 13778075) and grown for a total of 4 days, with passaging as cells reached high density.

### Drug treatments

Drugs were prepared as stock solutions at 1,000 to 10,000× the working concentration and diluted in growth medium before use. Drug name (abbreviation), solvent, working concentration, and source (catalog number) were as follows. Synthetic androgen R1881, ethanol, 2 nM, PerkinElmer (NLP005005MG). AHR ligand 10-Cl-BBQ (BBQ), DMSO, 100 nM, DC Chemicals (DC42712). RBN-2397 (RBN), DMSO, 10 to 100 nM, DC Chemicals (DC31069). FK866, ethanol, 2 to 10 nM, MedChemExpress (HY-50876). STF-118804 (STF), DMSO, 10 to 50 nM, MedChemExpress (HY-12808). Veliparib, DMSO, 1 μM, Selleck Chemicals (S100410MG). Bortezomib (Bort), DMSO, 1 μM, MedChemExpress (HY-10227). Cycloheximide (CHX), ethanol, 0.1 mg/ml, Alfa Aesar (AAJ6690103). Nicotinamide riboside (NR) was from Tocris Bioscience (6018) and was prepared as a 100× stock in sterile milliQ water at a working concentration of 1 mM. Drug-treatment durations are listed in the cell lines section.

For hypoxia experiments, cells were grown in a hypoxia chamber at 0.1% O_2_. Normoxic controls were grown under standard conditions at 20% O_2_.

### Total cell lysate preparation and Western blot analysis

Cells were grown and drug-treated in 12-well plates for the indicated durations. Growth medium was removed, and cells were optionally washed with PBS. SDS-PAGE loading buffer (200 mM Tris-HCl pH 6.8, 10% glycerol, 2% SDS, 3% β-mercaptoethanol; 0.12 ml/well) was added to the cells. The mixture was transferred to Eppendorf tubes, heated at 95°C for 5 minutes, cooled, sonicated (Branson Sonifier 250), and centrifuged at 16,000 × g for 3 minutes to obtain the supernatant. Proteins in the supernatant were separated by SDS-PAGE and transferred to nitrocellulose membranes. Membranes were blocked in 5% milk in PBST at room temperature for at least 45 minutes, incubated with primary antibody or fluorescently labeled FL-AF1521 probe in 1% BSA in PBST at room temperature for 2 hours or at 4°C overnight, washed five times in PBST (5 minutes each), incubated with fluorescently labeled secondary antibody in 1% milk in PBST at room temperature for 1.5 hours, and washed five times in PBST (5 minutes each). Signals were detected and quantified on an Odyssey CLx (LI-COR). Antibodies and probes are listed below.

| Antibody or probe | Source or reference | Clone # | WB usage |
| --- | --- | --- | --- |
| Mouse α-HA-tag | BioLegend / 901514 | 16b12 | 1:1,000 |
| Rabbit α-PARP7 (119-132 aa) | Yang et al, 2021 | | 1 μg/ml |
| ADPr probe FL-AF1521 | Kamata et al, 2019 | | 1 μg/ml |
| Mouse α-tubulin (TUB) | MilliporeSigma / T9028 | TUB-1A2 | 1:10,000 |
| mono-ADPr Fab antibody | Bio-Rad / TZA020 | AbD43647 | 1 μg/ml |
| Mouse α-DTX2 | ProteinTech / 67209-1-Ig | 2D4F8 | 1:3,000 |
| Rabbit α-DTX3L | Yang et al, 2017 | | 0.2 μg/ml |
| Rabbit α-AR (659-667 aa) | Yang et al, 2021 | | 1 μg/ml |
| Rabbit α-c-JUN | Cell Signaling / 9165S | 60A8 | 1:1,000 |
| Mouse α-FLAG-tag | MilliporeSigma / F3165 | M2 | 0.75 μg/ml |
| Mouse α-GST-tag | T. Parsons laboratory | mAb 9D9-F3-F7 hybridoma | 1 μg/ml |
| Mouse α-T7-tag | MilliporeSigma / 69522 | | 1:10,000 |
| Mouse α-lamin A/C (LMNA/C) | Abcam / ab40567 | Lamin A+C (JOL2) | 1:100 |
| Rabbit α-HIF1α | Cell Signaling / 3716 | | 1:1,000 |
| Rabbit α-C/EBPβ | Cell Signaling / 43095 | | 1:500 |
| AF680 donkey anti-mouse IgG (H+L) | Invitrogen / A10038 |  | 1:30,000 |
| AF680 donkey anti-rabbit IgG (H+L) | Invitrogen / A10043 |  | 1:30,000 |
| DyLight 800 goat anti-mouse IgG (H&L) | Rockland / 610-145-002 |  | 1:20,000 |
| DyLight 800 donkey anti-human IgG (H&L) | Invitrogen / SA5-10132 |  | 1:5,000 |

### Triton X-100 soluble and insoluble fractionation and protein pulldown

All steps were performed at 4°C. Harvested cell pellets were resuspended in Triton X-100 extraction buffer (20 mM Tris-HCl pH 7.5, 100 mM NaCl, 0.5% Triton X-100, 5 mM EDTA, 1 mM PMSF, 2 mM DTT, 5 μg/ml each of aprotinin, leupeptin, and pepstatin [A/L/P], and 1 μM veliparib) for 20 minutes and centrifuged at 18,000 × g for 20 minutes. The supernatant, defined as the Triton X-100 soluble fraction, was used for pulldown of most proteins in this study.

The remaining pellet (Triton X-100 insoluble fraction) was washed once or twice with extraction buffer, resuspended in 0.4 ml extraction buffer, sonicated (5 pulses at output 3 and 50% duty cycle, followed by a 1-minute pause, repeated four times), and centrifuged at 18,000 × g for 20 minutes. C/EBPβ was recovered in this supernatant.

Pulldowns were performed by incubating the supernatant with α-HA magnetic beads or with a GST-fusion protein immobilized on Glutathione MagBeads at 4°C for 3 to 4 hours. Beads were collected on a magnetic stand, and the supernatant was removed. Beads were washed five times in washing buffer (20 mM Tris-HCl pH 7.5, 100 mM NaCl, 0.1% Triton X-100, 1 mM EDTA, 2 mM DTT, 1 μg/ml each of A/L/P, and 1 μM veliparib), mixed with SDS-PAGE loading buffer, and heated at 95°C for 5 minutes.

For α-HA magnetic beads, 25 to 50 μl of bead slurry (Pierce 88836) was placed in an Eppendorf tube, collected on a magnetic stand, and washed with PBS containing 0.1% Triton X-100 after removing the supernatant. For GST-fusion beads, 10 μl of Glutathione MagBeads slurry (GenScript L00895-10) was mixed with 0.5 ml PBS containing 0.1% Triton X-100 in an Eppendorf tube, collected, and freed of supernatant, then incubated with 5 to 10 μg of GST-AF1521^Tandem^ (Kamata et al, 2019), GST-DTX2-RD, or GST-DTX2-RD^mut^ (Wierbiłowicz et al, 2025) in 0.2 ml PBS containing 0.1% Triton X-100 at 4°C overnight, then collected and washed.

### Cell permeabilization

Permeabilization was performed at 4°C. After growth medium was removed, cells in 12-well plates were washed once with 1 ml TBS (20 mM Tris-HCl pH 7.5, 100 mM NaCl) and permeabilized in 1 ml/well fractionation buffer (50 mM HEPES pH 7.5, 250 mM KCl, 2.5 mM MgCl_2_, 0.05% Triton X-100) for 20 minutes. The supernatant was removed, permeabilized cells were washed with 1 ml/well cold PBS, and 0.12 ml SDS-PAGE loading buffer was added to recover the chromatin-associated fractions.

### Protein half-life measurement

Protein half-life was measured in biological triplicate. Drug-pretreated cells were incubated with 0.1 mg/ml CHX in fresh growth medium for the indicated durations. Medium was removed immediately, and SDS-PAGE loading buffer was added to the cells. Samples were processed as for total cell lysate preparation. Proteins were separated by SDS-PAGE, transferred to nitrocellulose membrane, blotted with antibody or AF1521 probe, and detected and quantified on an Odyssey CLx (LI-COR). Signals were normalized to tubulin (TUB) and plotted in Prism 9 (GraphPad).

### Mercury treatment of ADP-ribosylated HA-PARP7

HA-PARP7-expressing cells were treated with 1 μM bortezomib for 4 hours before harvest. HA-PARP7 was extracted and immunoprecipitated on α-HA magnetic beads. Beads were washed twice with reducing agent-free buffer (100 mM Tris-HCl pH 7.5, 100 mM NaCl, 0.1% Triton X-100), aliquoted into individual tubes, and incubated at room temperature for 60 minutes in 100 mM Tris-HCl pH 7.5 containing 0 to 50 mM HgCl_2_, with MgCl_2_ added to maintain the divalent cation concentration where HgCl_2_ was reduced. Reactions were diluted with SDS-PAGE loading buffer and analyzed by SDS-PAGE and Western blot. ADP-ribose signals were normalized to HA signals.

### Ubiquitylation, NUDT16, and USP2-CD assays

HA-PARP7 was immunoprecipitated from cells pretreated with 1 μM bortezomib. After the immunoprecipitation wash, beads were washed once or twice with the respective assay buffer. Ubiquitylation assays were performed at 30°C for 30 minutes with HA-PARP7-bound beads in ubiquitylation buffer (20 mM Tris-HCl pH 7.5, 50 mM NaCl, 2 mM MgCl_2_, 1 mM DTT, 1 μg/ml each of A/L/P, 1 μM veliparib, 0.1 mg/ml BSA) supplemented with 1 mM ATP, 100 μg/ml T7-Ub (Yang et al, 2017), 5 μg/ml each of ubiquitin E1 (R&D Systems E-304) and ubiquitin E2 (His6-UbcH5C, R&D Systems E2-627), and 20 μg/ml GST-DTX2-RD or GST-DTX2-RD^mut^ (Wierbiłowicz et al, 2025). NUDT16 assays (NUDT16 from the In-Kwon Kim laboratory) were performed at 37°C for 60 minutes in NUDT16 buffer (20 mM Tris-HCl pH 7.5, 50 mM NaCl, 5 mM MgCl_2_, 2 mM DTT, 1 μg/ml each of A/L/P, 0.1 mg/ml BSA) supplemented with 0 to 25 μM NUDT16. USP2 assays were performed at 37°C for 15 minutes in USP2 buffer (50 mM Tris-HCl pH 7.5, 0.5 mM EDTA, 1 mM DTT, 1 μg/ml each of A/L/P, 0.1 mg/ml BSA) supplemented with 1 μM His-USP2-CD (catalytic domain, residues 258 to 606; Wierbiłowicz et al, 2025).

### Transcription assays

For transcription assays, cells were grown and drug-treated in phenol red-free RPMI 1640 (Gibco 11835-030) supplemented with 5% charcoal/dextran-stripped FBS (Gemini Bio-Products 100-119) and 1% P/S. RNA isolation (RNeasy kit, Qiagen 74104; QIAshredder, Qiagen 79656; DNase, Roche 04716728001), cDNA synthesis (iScript cDNA synthesis kit, Bio-Rad 178890), and qPCR (HotStart 2X Green qPCR Master Mix, APExBIO K1070) were performed according to the manufacturers’ instructions. qPCR primers were PARP7 (TCTCAGGAGCACTTGGAAAGA and TCAGCCTTCGTAGTTGGTCA), CYP1A1 (CAACCCTTCCCTGAATGCCT and GCTTCTCCTGACAGTGCTCA), and the GUS normalization control (CCGACTTCTCTGACAACCGACG and AGCCGACAAAATGCCGCAGACG).

### Total NAD and NAD^+^/NADH measurements

PC3-AR cells were treated with 100 nM BBQ and 2 nM FK866 for 18 hours, and total NAD and NADH were measured in technical triplicate using the WST method (Dojindo N509) with a 96-well plate reader (BioTek Synergy HTX) according to the manufacturer’s instructions. NAD^+^ was calculated by subtracting NADH from total NAD.

PC3-AR/HA-PARP7 cells were grown at 37°C in humidified incubators at either normoxic (21% O_2_/5% CO_2_/balance N_2_) or hypoxic (0.1% O_2_/5% CO_2_/balance N_2_; Avatar Odyssey incubator) conditions for 18 hours, with or without a final 40-minute treatment with 0.1 mg/ml CHX (administered under the appropriate O_2_ levels; inside a Coy Laboratory Glove Box for hypoxia). Cells were harvested under the appropriate O_2_ conditions, then NAD^+^ and total NAD were measured in technical triplicate for the normoxic and hypoxic conditions, or in technical duplicate for the hypoxic plus CHX condition, using a fluorometric method (Abcam ab176723, according to the manufactory’s instructions) in a 96-well plate reader (SpectraMax iD3). NADH levels were calculated by subtracting NAD^+^ from total NAD.

### Mass spectrometry

Mass spectrometric analysis was performed essentially as described previously (Hendriks *et al*, 2019) using quadruplicate biological replicates of PC3-AR cells treated with or without 2 nM R1881 for 15 h. Cells were washed twice with ice-cold PBS, harvested in PBS containing 1 μM olaparib, pelleted by centrifugation, lysed in 6 M guanidine-HCl (pH 8.5), snap-frozen, and stored at −80°C. Protein lysates were digested with Lysyl endopeptidase (Lys-C; 1:200, 3 h at room temperature), diluted threefold with 50 mM ammonium bicarbonate, and further digested with trypsin (1:100, overnight at room temperature). Peptides were purified using C18 Sep-Pak cartridges, lyophilized, and dissolved in AP buffer (50 mM Tris-HCl pH 8.0, 1 mM MgCl_2_, 250 μM DTT, and 50 mM NaCl). Potential ADP-ribose polymers were reduced to monomers by incubation with recombinant PARG (1:10,000, overnight at room temperature). ADP-ribosylated peptides were enriched using GST-tagged Af1521 macrodomain bound to Sepharose beads by incubation at 4°C for 3 h with head-over-tail rotation. Beads were washed with AP buffer, PBS, and water, and peptides were eluted with 0.15% TFA. Eluted peptides were basified and fractionated into three fractions using C18 StageTips prior to LC-MS/MS analysis.

LC-MS/MS analysis was performed using an EASY-nLC 1200 system coupled to an Orbitrap Fusion Lumos mass spectrometer (Thermo Fisher Scientific). Peptides were separated on a 15 cm × 75 μm i.d. analytical column packed in-house with 1.9 μm C18 beads using a 50 min gradient from 3% to 38% buffer B (80% acetonitrile, 0.1% formic acid). The mass spectrometer was operated in data-dependent acquisition mode with full MS scans acquired at a resolution of 120,000 and a maximum injection time of 50 ms. Precursors with charge states 3-5 were fragmented using EThcD with supplemental activation energy set to 20. MS/MS spectra were acquired in the Orbitrap at a resolution of 60,000 with a maximum injection time of 500 ms.

Raw files were analyzed using MaxQuant (v1.5.3.30) and searched against the human UniProt database (downloaded April 29, 2023). Carbamidomethylation of cysteine and ADP-ribosylation of C, D, E, H, K, R, S, T, and Y residues were included as variable modifications. A maximum of five variable modifications and four missed cleavages were allowed. The minimum Andromeda delta score for ADP-ribosylated peptides was set to 20.

### Data availability

The mass spectrometry proteomics data have been deposited to the ProteomeXchange Consortium via the PRIDE partner repository (Perez-Riverol *et al*, 2025) with the dataset identifier PXD084367.

### Reviewer access details

Log in to the PRIDE website using the following details: Project accession: PXD084367, Token: vDvhRXBjmDlI

### NAD^+^ docking against PARP7

Molecular docking of NAD^+^ to PARP7 was performed with AutoDock Vina v1.2.6(Eberhardt *et al*, 2021; Trott & Olson, 2010). The PARP7 structure was obtained from the AlphaFold Protein Structure Database (model AF-Q7Z3E1-F1)(Jumper *et al*, 2021; Varadi *et al*, 2022) and converted to PDBQT format with ADFRsuite v1.0 prepare_receptor (Ravindranath *et al*, 2015), adding hydrogens with the -A hydrogens flag. The NAD^+^ SMILES string (NC(=O)c1ccc[n+]([C@@H]2O[C@H](COP(=O)(O)OP(=O)(O)OC[C@H]3O[C@@H](n4cnc5c(N)ncnc54)[C@H](O)[C@@H]3O)[C@@H](O)[C@H]2O)c1) was converted to PDBQT format with RDKit v2025.09.5 by adding hydrogens, generating 3D coordinates with the ETKDGv3 conformer-embedding algorithm, and minimizing energy with the MMFF94s force field, followed by conversion with Meeko v0.7.1. Docking used a box of 22 × 22 × 22 Å and an exhaustiveness of 64. All other parameters were AutoDock Vina defaults.

### Bioinformatics analysis of tumor and cell line datasets

TCGA-PRAD RNA-seq data (HiSeqV2; log (norm_count + 1)) were downloaded from the UCSC Xena platform (Goldman *et al*, 2020) and restricted to primary tumor samples (code “01”; n = 497)(Goldman *et al*, 2020). Molecular annotation (tumor purity, reviewed Gleason sum, AR score, molecular subtype) were downloaded from Abeshouse *et al*. (Abeshouse *et al*, 2015) Supplementary Table S1A, and the survival endpoints were downloaded from TCGA Clinical Data Resource (Liu *et al*, 2018), with progression-free interval (PFI) as the primary endpoint. We obtained DDR/genomic instability scores (HRD score = TAI + LST + LOH; BRCA mutational signature, COSMIC Signature 3; aneuploidy score) from Knijnenburg *et al*. (Knijnenburg *et al*, 2018), filtered to PRAD (n = 464–479).

SU2C/PCF 2019 mCRPC data (Abida *et al*, 2019) were accessed via cBioPortal. Fifteen of 429 patients had two biopsies profiled, so we retained one sample per patient using the alphabetically first SAMPLE_ID, giving n = 262. Cell line expression (log (TPM + 1)) and RBN2397 viability (PRISM Repurposing Primary, log fold change) were downloaded from DepMap (Corsello *et al*, 2020) release 26Q1.

We scored samples for hypoxia using ssGSEA (GSVA ≥ 1.44; ssgseaParam, normalize = TRUE, minSize = 5) with the MSigDB HALLMARK_HYPOXIA gene set (msigdbr), computed independently per cohort. All correlations presented in the reported bioinformatic analyses are Spearman rank correlations, with p-values corrected by Benjamini-Hochberg within each analysis panel. Partial Spearman correlations were computed by rank-transforming variables, regressing covariates from both variables by linear regression on ranks, and correlating the residuals. We tested predictor-by-subtype and predictor-by-moderator interactions by comparing a rank-based linear model with and without a continuous interaction term by F-test. Group comparisons used the Wilcoxon rank-sum test (two groups), Kruskal-Wallis test (three or more groups), or Fisher’s exact test for categorical associations.

Cox regression (coxph, survival package) was used for PFI with z-scored predictors across three nested covariate tiers: (1) Gleason grade and AR score; (Sramkoski *et al*) age, T stage, and Gleason grade; and (3) tier 2 plus molecular subtype. The 12-resulting p-values were corrected jointly by Benjamini-Hochberg. We performed a quartile analysis (top vs. bottom 25%, age-adjusted Cox) to replicate Yang et al. (2021). Kaplan-Meier curves (survminer, median split) are for visualization only.

For AP-1 gene set enrichment analysis, RNA-seq data from VCaP cells treated for 18 hours with vehicle, R1881 (2 nM), RBN-2397 (100 nM), or both compounds (three biological replicates per condition) were analyzed using DESeq2 (v1.50.1). Genes with at least 10 counts in three or more samples were retained for analysis. Differential expression was assessed using the Wald test, and genes were ranked by their Wald statistic for the R1881 + RBN-2397 versus R1881 comparison. Gene set enrichment analysis (GSEA) was performed using seven experimentally derived AP-1 family transcription factor target gene sets from the ChEA 2022 library (Chen *et al*, 2013; Lachmann *et al*, 2010), obtained through Enrichr (https://maayanlab.cloud/Enrichr/). Enrichment was assessed using the fgsea package, with P-values adjusted for multiple testing using the Benjamini-Hochberg method across the seven gene sets.

For AHR-dependent gene expression analysis, RNA-seq data from VCaP cells treated for 18 hours with vehicle, BBQ (100 nM), RBN-2397 (100 nM), or both compounds (three biological replicates per condition) were analyzed using DESeq2 (v1.50.1). Genes with at least 10 counts in three or more samples were retained. Differential expression was assessed using a factorial model including BBQ, RBN-2397, and their interaction. The interaction coefficient represents the effect of RBN-2397 in BBQ-treated cells minus its effect in untreated cells. Genes with Benjamini-Hochberg-adjusted P < 0.05 for the interaction term were considered significant.

All analyses were performed in R version 4.4.1 or 4.5.0 using tidyverse, readxl, msigdbr, GSVA, ggpubr, ggcorrplot, survival, survminer, DESeq2, fgsea, and related Bioconductor packages.

### Software and statistical analysis

Protein half-life, qPCR, and related assay data were plotted in Prism 9 (GraphPad), except for the fluorometric NAD^+^/NADH standard-curve fit (Abcam ab176723), which was performed in Excel (Microsoft). Error bars represent standard deviation. Significance (P) for these assays was determined by two-tailed unpaired t test (**** P < 0.0001, *** P < 0.001, ** P < 0.01). Statistical analysis of the tumor and cell line datasets is described in the preceding section.

## FIGURE LEGENDS

**Supplementary Fig 1.**
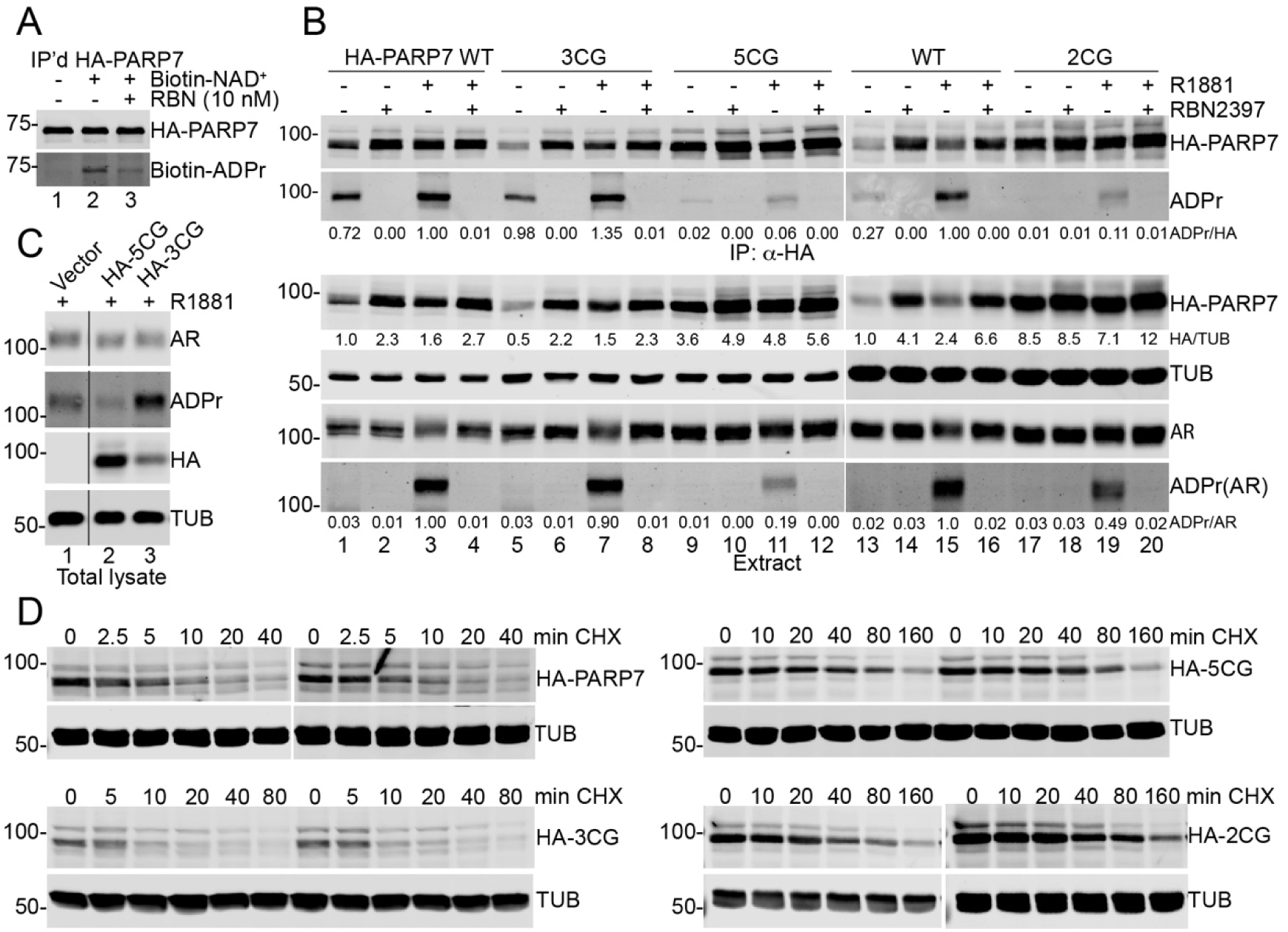
ADP-ribosylation site mutations affect PARP7 activity and stability. **A.** Auto-ADP-ribosylation of IP’d HA-PARP7 and its inhibition by RBN-2397. **B.** C39,99,439G (3CG) mutation does not affect PARP7 activity and stability while C543, 552G mutation increases PARP7 protein stability and reduces its ADP-ribosylation activity toward AR or itself. Cell extracts and IP’d HA-PARP7 wild type or mutants were subjected to antibody or FL-AF1521 detection. **C.** AR ADP-ribosylation from the HA-PARP7 5CG mutant (all 5 sites) is from the endogenous PARP7. **D.** HA-PARP7 WT and mutants’ blotting data for half-life measurements related to Figure 2E.

**Supplementary Fig 2.**
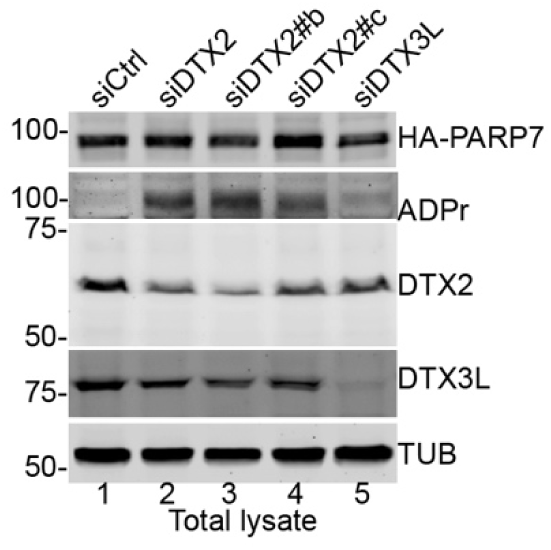
PARP7 ADP-ribosylation was affected by knockdown of DTX2 but not DTX3L. PC3-m cells that stably transfected with HA-PARP7 were transiently transfected with siCTRL (Ambion AM463S), SÌDTX2 (Ambion S223231), siDTX2#b (Ambion S41519), s-DTX2#c (Ambion S195603) or siDTX3L (target sequence CGCGTATTAGGAGTCTCAGAT) for 4 days. Total cell lysates were subjected to SDS-PAGE. Proteins and ADPr were detected using antibodies against HA, DTX2, DTX3L, Tubulin (TUB) and FL-Af1521.

**Supplementary Fig. 3.**
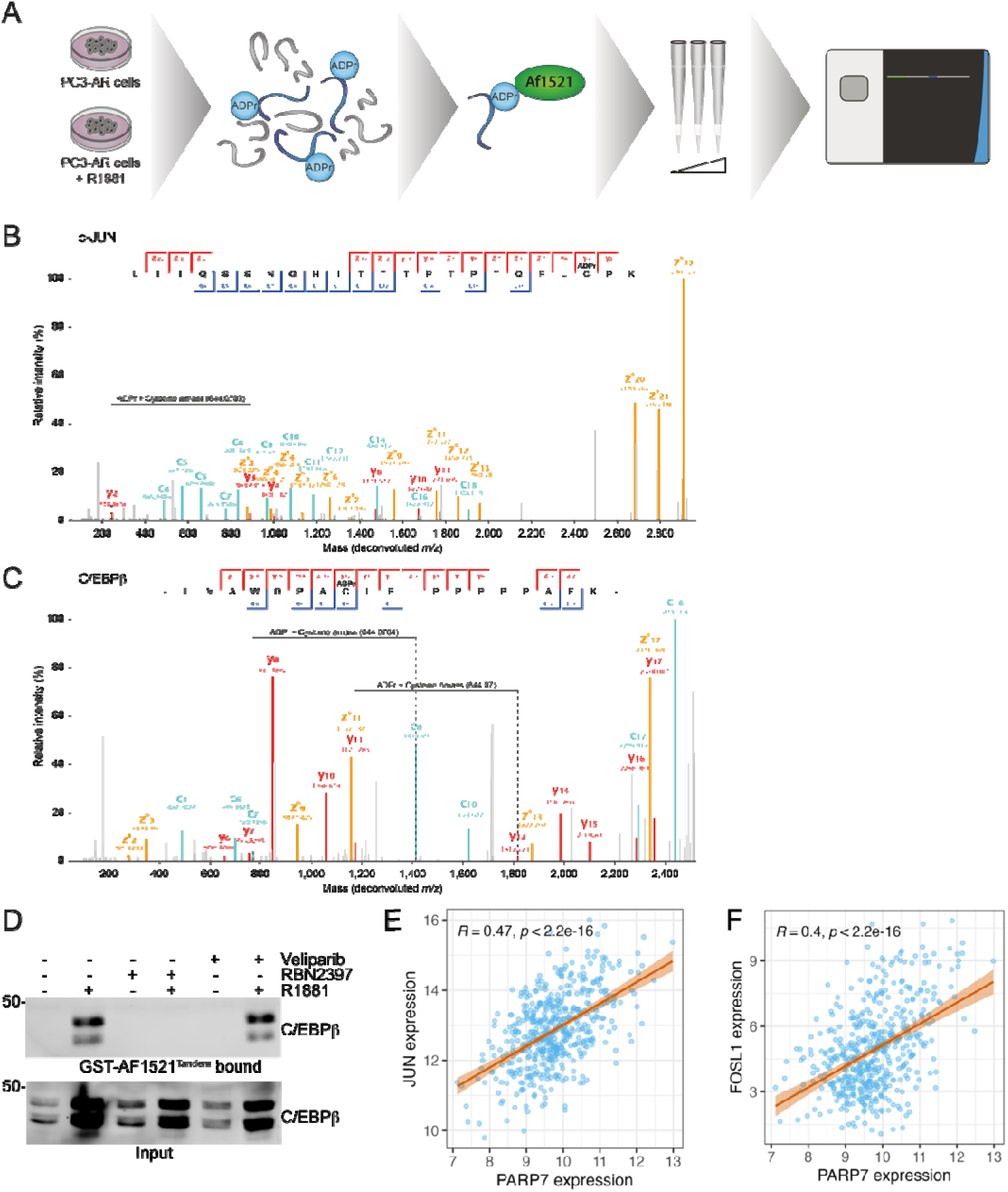
PARP7 ADP-ribosylates c-JUN and C/EBPβ, and PARP7 mRNA correlates with JUN and FOSL1 expression in prostate cancer. **A.** Workflow illustrating the experimental setup. Control cells and cells treated with R1881 were prepared in quadruplicate, lysed, digested, and enriched using the AÍ1521 macrodomain. ADPr enriched peptides were analyzed using EThcD on an Orbitrap Fusion Lumos. **B.** Fully annotated MS/MS spectrum corresponding to a peptide from o-JUN modified by ADPr on C-9B. **C.** Fully annotated MS/MS spectrum corresponding to a peptide from C/EBPB modified by ADPr on C-11. **D.** C/EBPβ was ADP-riboaylated In the presence of high PARP7 level when PC3AR cells were androgen-treated; the ADP-ribo- βylatlon was Inhibited by a PARP7 inhibitor RBN-2397 but not a PARP1/2 inhibitor Vellparib. PC3-AR cells were treated with 2nM R1881, lOOnM rbn and lOOnM vellparib tor 17 hrs, harvested and Triton x-100 extracted. The insoluble fractions wore further treated with sonication followed with centrifugation (see methods section). These soluble fractions (also inputs) were subjected to GST-AflSZI™“*” beads pulldown. Inputs and beads bound fractions were separated on SDS-PAGE followed with Western blot analyses using C/EBPβ antibody. **E.** PARP7 expression versus JUN expression in TCGA-PRAD primary tumors (n=496). Expression values are log_2_-transformed normalized counts. Spearman p and p-value annotated; regression line with 95% confidence Interval shown. **F.** PARP7 expression versus FOSL1 expression in TCGA-PRAD primary tumors (n=49θ). Expression values are log_a_-tranβ- formed normalized counts. Spearman p and p-value annotated; regression line with 95% confidence Interval shown.

**Supplementary Fig 4.**
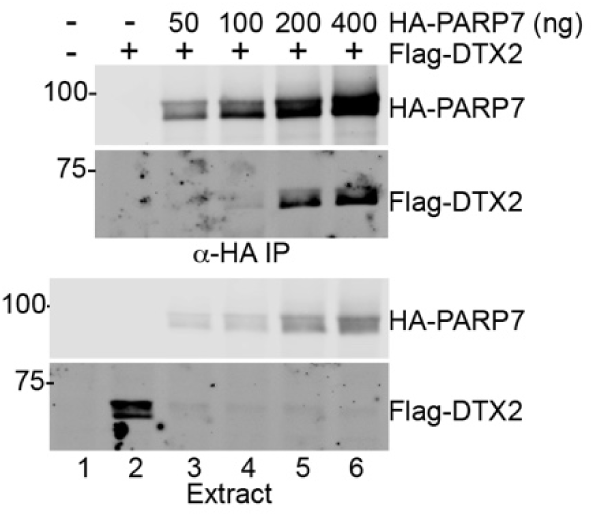
PARP7 interacts with DTX2. The pcDNA3/Flag-DTX2 plasmid DNA (2 pg) and 0.4 pg of combined plasmid DNA of pKH3/HA-PARP7 and vector were co-transfected into 60 mm plates of HEK293T cells for 20 hrs. Cells were incubated with 1 uM of Bortezomib in the fresh medium for 4 hrs before cell harvest. Cell extracts were subjected to anti-HA immunoprecitation. Proteins were separated on SDS-PAGE and detected using anti-HA and M2 Flag-tag antibodies.

**Supplementary Fig 5.**
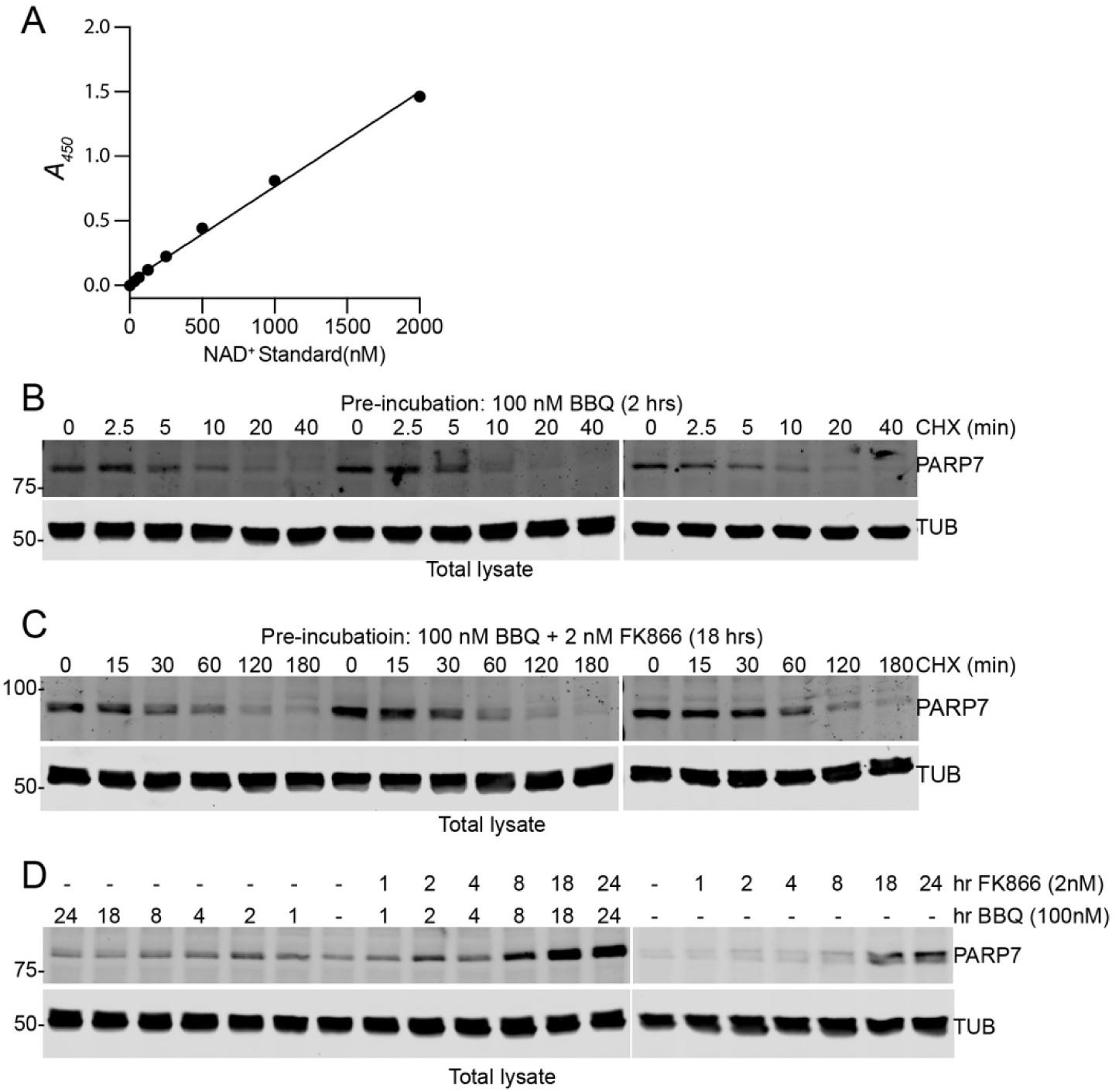
NAMPT inhibitor FK866 stabilizes PARP7 protein. **A.** NAD^+^ standards using the NAD/NADH Assay Kit - WST. **B.** Cycloheximide (CHX) chase experiments to measure PARP7 half-life after PC3AR cells were pre-treated 2 hrs with 100nM BBQ. **C.** Cycloheximide (CHX) chase experiments to measure PARP7 half-life after PC3AR cells were pre-treated 18 hrs with 100nM BBQ and 2nM FK866. **D.** PARP7 protein level at the indicated timepoints with BBQ or RBN alone or combination.

**Supplementary Fig 6.**
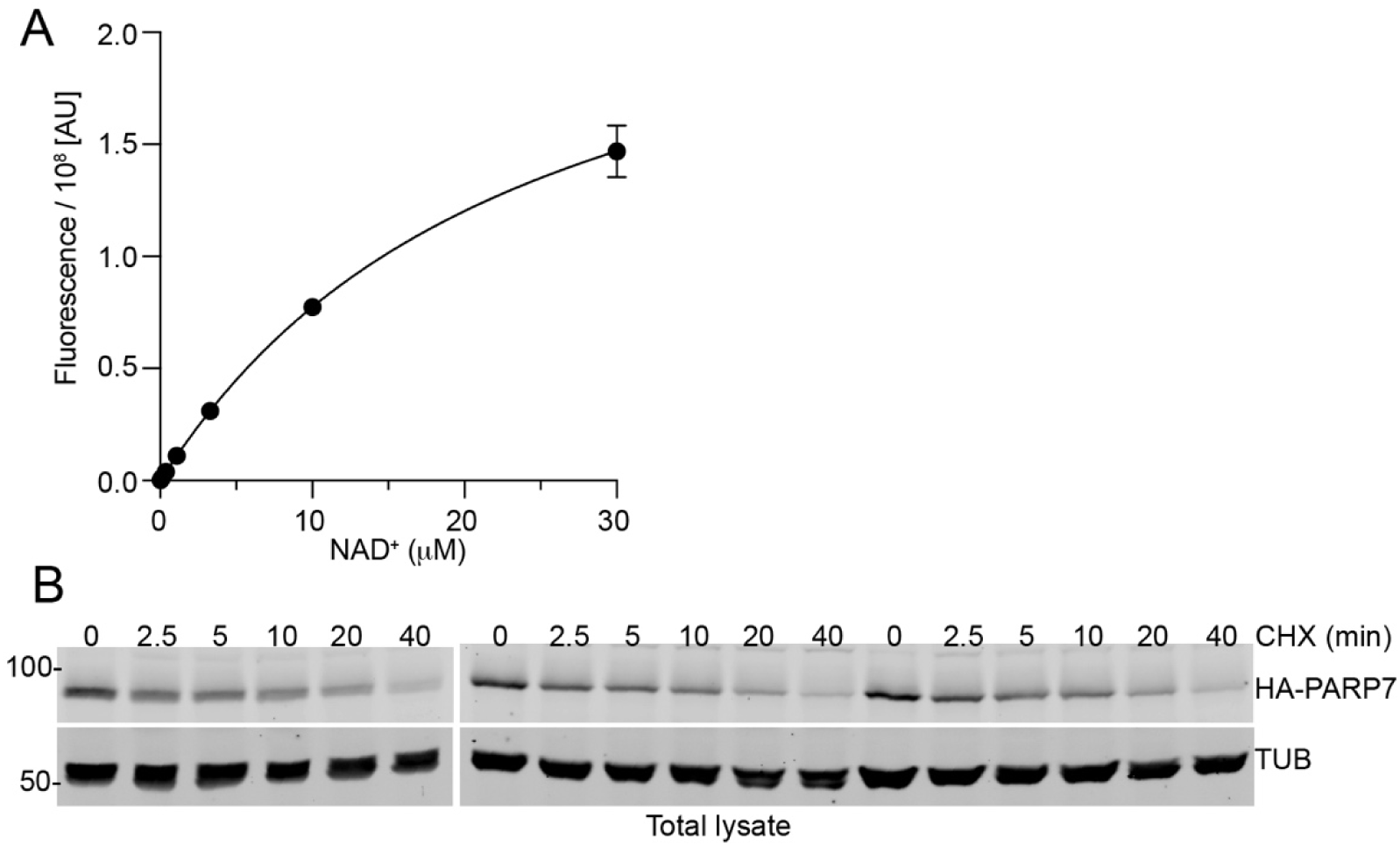
Hypoxia condition stabilizes PARP7 protein. **A**.NAD^+^ standards using the NAD/NADH Assay Kit (Fluorometric). **B**. Cycloheximide (CHX) chase experiments to measure PARP7 half-life after PC3AR(HA-PARP7) cells were under hypoxia condition (0.1% O_2_) for 18 hrs.

**Supplementary Fig 7.**
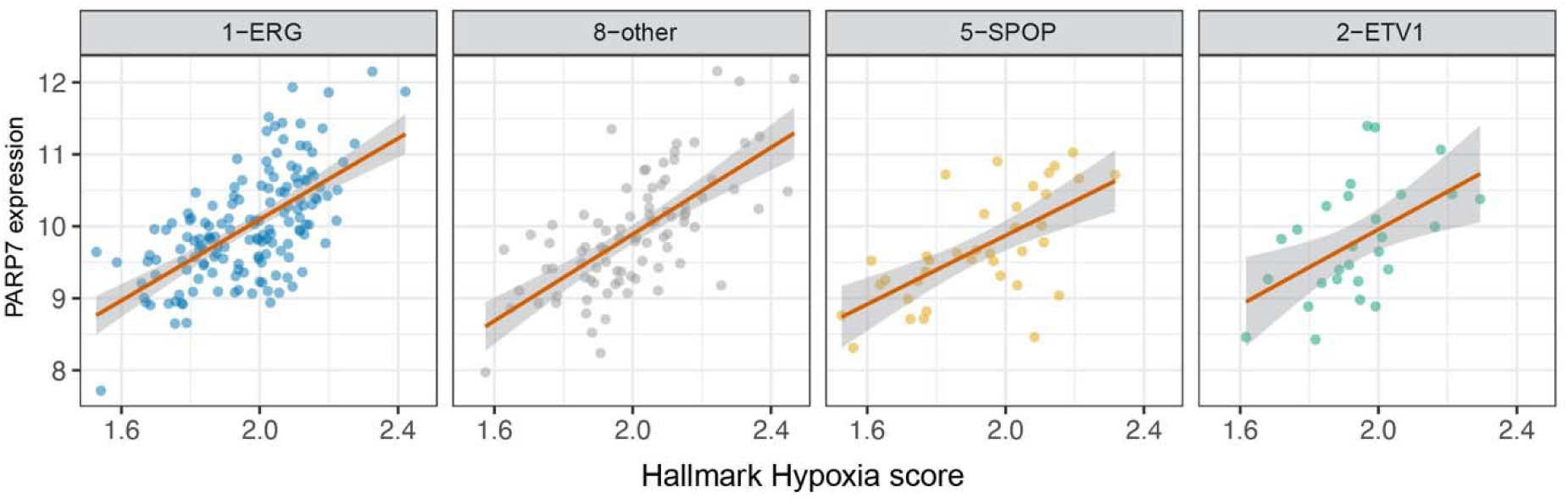
Hallmark Hypoxia versus PARP7 by molecular subtype in TCGA-PRAD. Hallmark Hypoxia ssGSEA score versus PARP7 expression stratified by TCGA-PRAD molecular subtype (ERG-fusion, ETV1-fusion, SPOP-mutant, unclassified; subtypes with n≥20 retained). Expression values are log_2_-transformed normalized counts. Regression lines fitted independently per subtype.

